# Winter in water: Differential responses and the maintenance of biodiversity

**DOI:** 10.1101/849109

**Authors:** Bailey C. McMeans, Kevin S. McCann, Matthew M. Guzzo, Timothy J. Bartley, Carling Bieg, Paul Blanchfield, Timothy Fernandes, Henrique Giacomini, Trevor Middel, Michael Rennie, Mark Ridgway, Brian Shuter

## Abstract

The ecological consequences of winter in freshwater systems are an understudied but rapidly emerging research area. Here, we argue that winter periods of reduced temperature and light (and potentially oxygen and resources) could play an underappreciated role in mediating the coexistence of species. This may be especially true for temperate and subarctic lakes, where seasonal changes in the thermal environment might fundamentally structure species interactions. With climate change already shortening ice-covered periods on temperate and polar lakes, consideration of how winter conditions shape biotic interactions is urgently needed. Using freshwater fishes in northern temperate lakes as a case study, we demonstrate how physiological trait differences (e.g., thermal preference, light sensitivity) drive differential responses to winter among competing species. Specifically, some species have higher a capacity for winter activity than others. Existing and new theory is presented to argue that such differential responses to winter can promote species coexistence. Importantly, if winter is a driver of niche differences that weaken competition among species, then shrinking winter periods could threaten coexistence by tipping the scales in favor of certain sets of species over others.

## Introduction

The role of temporal variation for promoting species coexistence has a long history in ecology (e.g., Paradox of the Plankton, Hutchinson 1961). Just as species select for different resources and habitats, and in doing so partition their niche in space, species can also diverge in the timing of their activity, partitioning their niche in time (Chesson 2000). Modern coexistence theory asserts that temporal niche divergence in fluctuating environments can promote coexistence that would otherwise be impossible in a static environment (Adler et al. 2006; Angert et al. 2009; Chesson & Huntley 1997; Tredennick et al. 2017).While much of this existing theory has focused on inter-annual variation, seasonal variation is being increasingly recognized for its role in coexistence (Mathias & Chesson 2013; Mellard et al. 2019; Shimadzu et al. 2013; Tonkin et al. 2017; Treddnick et al. 2017). For example, competing species inhabiting the same environment and experiencing the same conditions can coexist by diverging in their annual patterns of metabolic activity (Szabo et al. 2016).

The onset and retreat of winter brings pronounced temperature variation across temperate and polar latitudes, with many lakes becoming ice-covered during winter. In these regions, winter in water is characterized by annual temperature minimums throughout the water column (0 to 4°C), reduced light, and potentially reduced oxygen and resource density (Shuter et al. 2012). Biota respond to these prominent seasonal changes in abiotic conditions in a variety of ways, including foraging, growing and reproducing at different times of the year. Because the metabolic rate of ectotherms is directly related to temperature, winter temperatures should ubiquitously suppress the ability of fish and other aquatic organisms to move, capture and digest prey, avoid predators, and grow (Hurst 2007). Yet, many taxonomic groups, including amphipods (Werner 2006), zooplankton (Mariash et al. 2017) and fish (Shuter et al. 2012) contain species that remain active during winter. Some fishes, for example, are capable of maintaining or even gaining biomass under ice cover, owing to physiological adaptations that facilitate foraging in dark, cold conditions (Fig. 1A; Byström et al. 2006; French et al. 2014). Fishes that perform best at colder temperatures (i.e. with colder thermal preferences) are expected to have a higher capacity for winter activity than fishes with warmer thermal preferences (Fig. 1B). Fishes that lack adaptations to low light and cold should be inefficient foragers during the winter, and are expected to adopt an overwintering strategy of suppressed activity (Fig. 1B; Watson et al. 2019). Because all ectotherms should be active and growing during warmer, brighter, open water periods (but within their upper thermal limit), divergent responses to cold, dark winters could be a mechanism for promoting niche partitioning and thermal performance trade-offs that favor different species at different times (Fig. 1C). Such trade-offs can promote coexistence (Angert et al. 2011). However, winter is historically understudied in ecology, and consideration of how cold, ice-covered periods shape species interactions in aquatic ecosystems is in its infancy (Helland et al. 2011; Salonen et al. 2009).

**Figure 1.**
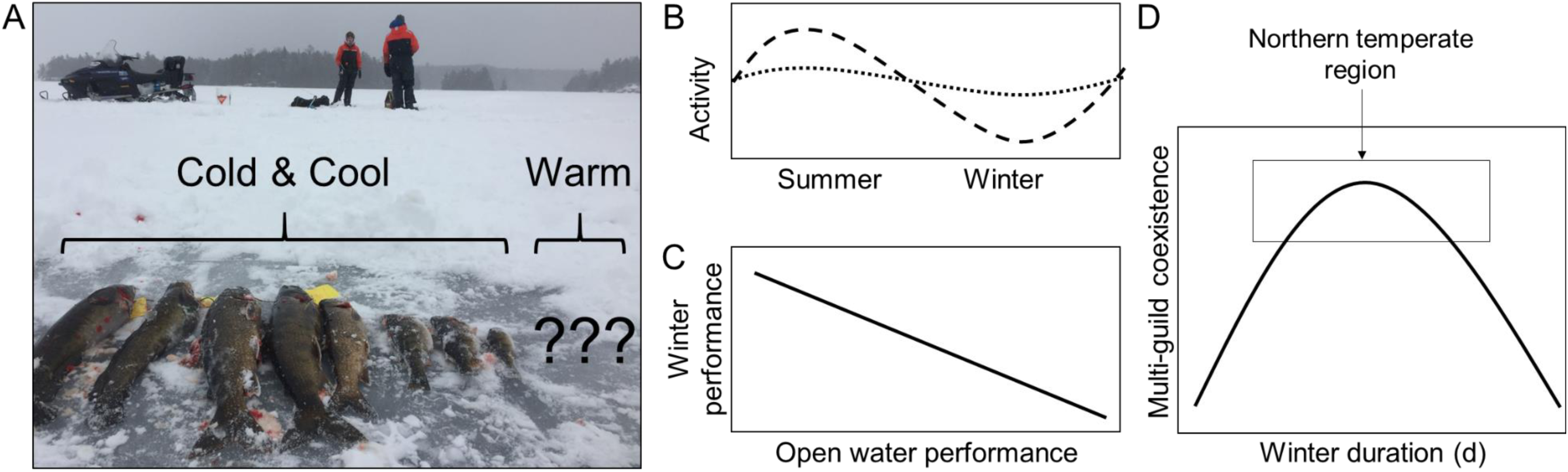
A) Northern temperate freshwater fishes vary widely in their responses to winter. Those that are better adapted for winter activity are easily captured during winter sampling operations and tend to have colder thermal preferences (A). Other species can be more challenging to capture under ice cover and tend to have warmer thermal preferences (e.g., bass and other sunfishes; A). Being ectotherms, all fishes should be actively foraging during warmer, brighter open water periods, and have reduced activity rates during winter (B). Fishes with colder thermal preferences (short dash), however, are expected to have a higher capacity for winter activity than species with warmer thermal preferences (long dash; B). Optimal performance under a particular set of conditions implies reduced performance under other conditions and sets up a variety of possible trade-offs (Angert et al. 2009; Kingsolver 2009). For example, a high capacity for activity during warm, bright, open water months is expected to trade-off with a reduced capacity for activity during cold, dark winters (C). Seasonal variation broadly and winter periods specifically could therefore favor different species at different times and promote coexistence of species belonging to multiple thermal guilds (D). In northern temperate lakes, winters are likely long enough to support cold-adapted species but not too long so as to exclude warm-adapted species, making this **winter-mediated species coexistence** particularly likely (D).

Here, we argue that the conditions imposed by winter and the different adaptations of species to succeed under such conditions could play an underappreciated role in species coexistence. This **winter-mediated species coexistence** could be widespread, but is especially probable in northern temperate regions that experience winters of intermediate duration (Fig. 1D). We first draw from modern theory to discuss how winter conditions could promote coexistence. We then use new and existing empirical data to demonstrate how freshwater fishes inhabiting northern temperate lakes diverge in their response to winter in ways that are consistent with previously described coexistence mechanisms. Next, we make the case that species likely diverge in time and space simultaneously, by both being active at different times and by partitioning habitats and resources in space. Finally, we present novel theory to explore when and where winter is most important for coexistence, and we pose these ideas as testable predictions for future work. Although winter is broadly viewed as a stressful period that limits ectotherm survival and growth, our perspective is that seasonal variation (and differential responses to winter periods, specifically), could also shape biotic interactions in ways that maintain diversity.

### Coexistence in fluctuating environments

The ecological implications of temporally fluctuating conditions have intrigued ecologists for decades (Hutchinson 1961; Schoener 1982; Wiens 1977). Based on modern coexistence theory, non-equilibrium conditions alone do not promote coexistence simply by reducing population growth rates or decreasing the importance of competition (Chesson & Huntley 1997; Hart & Marshall 2013). Instead, to have a positive influence on coexistence, fluctuating conditions must either reduce the population growth rate of the dominant species (i.e., equalizing mechanisms) or generate opportunities for niche differentiation in space or time that weaken the strength of inter- vs. intraspecific competition (i.e., stabilizing mechanisms; Chesson 2000, 2018).

#### Equalizing mechanisms

Harsh periods that reduce population growth rates and amplify population fluctuations can increase sensitivity to competition and lead to extinction and diversity loss (Chesson & Huntley 1997). However, harsh periods can also reduce differences in species average fitness if they disproportionately affect the dominant species (i.e., the equalizing mechanism; Chesson 2018; Chesson & Huntley 1997). Here, the term ‘species’ in ‘species average fitness’ differentiates the concept from individual fitness, and ‘average’ refers to the average both: 1) across all individuals within a population of a particular species, and 2) across all environmental conditions experienced by that population (Chesson 2018). Species with high average fitness have a high growth rate and/or a low sensitivity to competition, giving them a competitive advantage over species with lower average fitness (Chesson 2000, 2018). Harsh periods, including but not restricted to winter, can operate on coexistence as an equalizing mechanism that reduces differences in species average fitness, thus slowing competitive exclusion and allowing otherwise competitively inferior species to persist for longer (Chesson & Huntley 1997).

Importantly, while equalizing mechanisms can dampen the competitive edge of the dominant species, they do not singly guarantee stable coexistence. Instead, the species with the higher average fitness will eventually win and exclude the species with lower average fitness. Even if species have identical average fitness, neutral theory predicts they will eventually drift towards competitive exclusion, whose probability and/or timing would depend on population sizes and turnover rates (Hubbell 2011). Stabilizing mechanisms are required to overcome ecological drift to extinction or any differences in species average fitness (Adler et al. 2007; Chesson 2018; Tredennick et al. 2017).

#### Stabilizing mechanisms

Species may keep ‘out of each other’s way’ by using different habitats or by partitioning resources in ways that do not depend on temporal fluctuations (Chesson 2000). So-called fluctuation-independent mechanisms instead rely on spatial heterogeneity (Chase & Leibold 2002; Macarthur & Levins 1964). Similar to resource partitioning in space, species can differentiate in the timing of when they use a shared resource (e.g. by feeding, growing, and reproducing at different times; Chesson 1985). Two ‘fluctuation-dependent’ coexistence mechanisms are recognized: relative non-linearity and the storage effect (Chesson 2018). Relative non-linearity is when species differ in the shape of their functional response curves (i.e., one species has a more non-linear response to changes in resource density than the other species), favoring different species under different resource conditions (Armstrong & McGehee 1980). The storage effect operates when species are able to ‘store’ during good times to sustain positive population growth during bad times. Both relative non-linearity and the storage effect operate via fluctuations (in resources or environmental conditions) that favor different species at different times, allowing coexistence that would be impossible in a static environment (Chesson 2018).

Harshness can act as a stabilizing mechanism when it provides opportunities for niche partitioning (Chesson & Huntley 1997). Niche partitioning in time, just as in resource partitioning in space, weakens the strength of inter-specific competition (vs. intra-specific competition) and allows an inferior competitor or low-density species to coexist with a superior competitor. Empirical evidence supports both relative non-linearity and the storage effect as coexistence mechanisms operating in nature, albeit mostly for plants and plankton (Adler et al. 2006; Angert et al. 2011; Caceres 1997; Chesson et al. 2012; Descamps-Julien &Gonzalez 2005; Zepeda & Martorell 2019). Relative non-linearity could contribute most strongly to coexistence in species that diverge in life-history traits (e.g., fast vs. slow strategies) but otherwise have high niche overlap (Xiao & Fussmann 2013). For the storage effect, differential responses to the environment (condition #1 of the storage effect) tend to generate environment-competition covariance (condition #2), and species with long-lived adults have buffered population growth (condition #3; Chesson et al. 2012). Northern temperate freshwater fishes are known to vary in their life-history traits (e.g., spawn time, age at maturity, fecundity, lifespan) and span broad gradients in optimal environmental conditions for growth and metabolism (e.g. temperature; King et al. 1999). Fluctuation-dependent coexistence mechanisms could therefore operate in these communities.

### Winter in water: differential responses and the maintenance of biodiversity

#### Winter in northern temperate lakes

Temperate latitudes are characterized by four distinct seasons. For waterbodies in this region, winter has been previously defined as the period of ice cover (Shuter et al. 2012). Southern temperate lakes below about 40°N do not experience stable ice cover, so do not have ‘tru’ winters, based on this definition. Lakes above about 60°N in subarctic and Arctic zones, on the other hand, experience long winters that can last for over half of the year, which can restrict biodiversity to only the most cold-adapted fishes. Northern temperate lakes—from about 40 to 60°N—fall between these two extremes, with ice cover lasting anywhere from 1 to 6 months depending on geographic location and elevation (Shuter et al. 2012). Northern temperate winters are therefore short enough to allow a phylogenetically and physiologically diverse fish assemblage to establish (unlike Arctic lakes), while also experiencing a unique set of conditions associated with stable ice cover (unlike southern temperate lakes; Shuter & Post 1990).

Lakes <61°N are also experiencing rapid change due to climate warming and are at the highest risk of ice cover loss (Weyhenmeyer et al. 2011). Currently, most northern temperate lakes are dimictic, meaning warm surface waters stratify from cooler water at depth during summer, cold temperatures establish throughout the water column during winter, and the lake mixes twice per year (fall and spring). Warming temperatures are threatening to shift northern temperate lakes from ice-covered and dimictic to ice-free and monomictic (Weyhenmeyer et al. 2011). This is a concern because warmer and more consistently stratified water columns will restrict cold-adapted fishes from accessing nearshore habitats and prey, with negative consequences for their growth (Guzzo et al. 2016; Plumb et al. 2014).

Along with cold temperatures, winter brings a shorter photoperiod that reduces light availability in the underlying water column. Ice and especially snow cover further attenuate the light available for primary production (Jewson et al. 2009) and foraging for visual predators (Blanchfield et al. 2009). Ice cover also restricts atmospheric exchange, which can produce low oxygen conditions (Terzhevik et al. 2009) that affect fish survival, and therefore community structure (Hurst 2007; Tonn & Magnuson 1982). Resources available for fishes could also change seasonally. Overwinter mortality of small-bodied minnows, for example, could reduce prey density for piscivores (Rennie et al. 2019), although very few studies have directly quantified prey density during ice cover. Even if resources remain at comparable densities between open water and ice cover, a particular resource may not be accessible during winter (e.g., due to low light; Blanchfield et al. 2009). However, some fishes are more tolerant of winter conditions than others, remaining active and successfully foraging despite the cold and dark conditions.

#### Differential responses to winter in fish

Temperature is a fundamental niche axis for ectotherms that dictates all major metabolic processes (Fry 1947). Fishes, like other ectotherms, have a temperature range over which they perform optimally with respect to their activity, growth and reproduction (Casselman 2002; Hokanson 1977; Pörtner 2007). Temperature preferences of freshwater fishes fall along a continuous spectrum, but three discrete categories are typically used based on their optimal thermal performance temperatures: the cold-water, the cool-water and warm-water guilds (Fig. 2A; Casselman 2002; Hokanson 1977; Magnuson et al. 1979). Fig. 2A illustrates the guild membership, temperature preference and upper thermal limit for a sample of common North American freshwater fishes, using typical threshold values for each guild (<17.5°C for cold-water, 17.5 to <25°C for cool-water, and >25°C for warm-water). Thermal preferences reflect the adaptation of: 1) molecular processes (e.g., stability vs. flexibility of proteins, cell membranes, DNA and RNA; Pörtner 2007), and 2) capacity to supply oxygen at the whole organism level (e.g., aerobic scope; Pörtner 2002) to a particular temperature range. Because winter temperatures of 4°C are below the thermal optima of nearly all North America freshwater fishes (Fig. 2A), such temperatures should generally reduce the aerobic scope of all species. As a result, available thermal habitat to achieve optimal growth is typically plentiful for all fish of all thermal guilds at some time during open water, but absent during winter.

**Figure 2.**
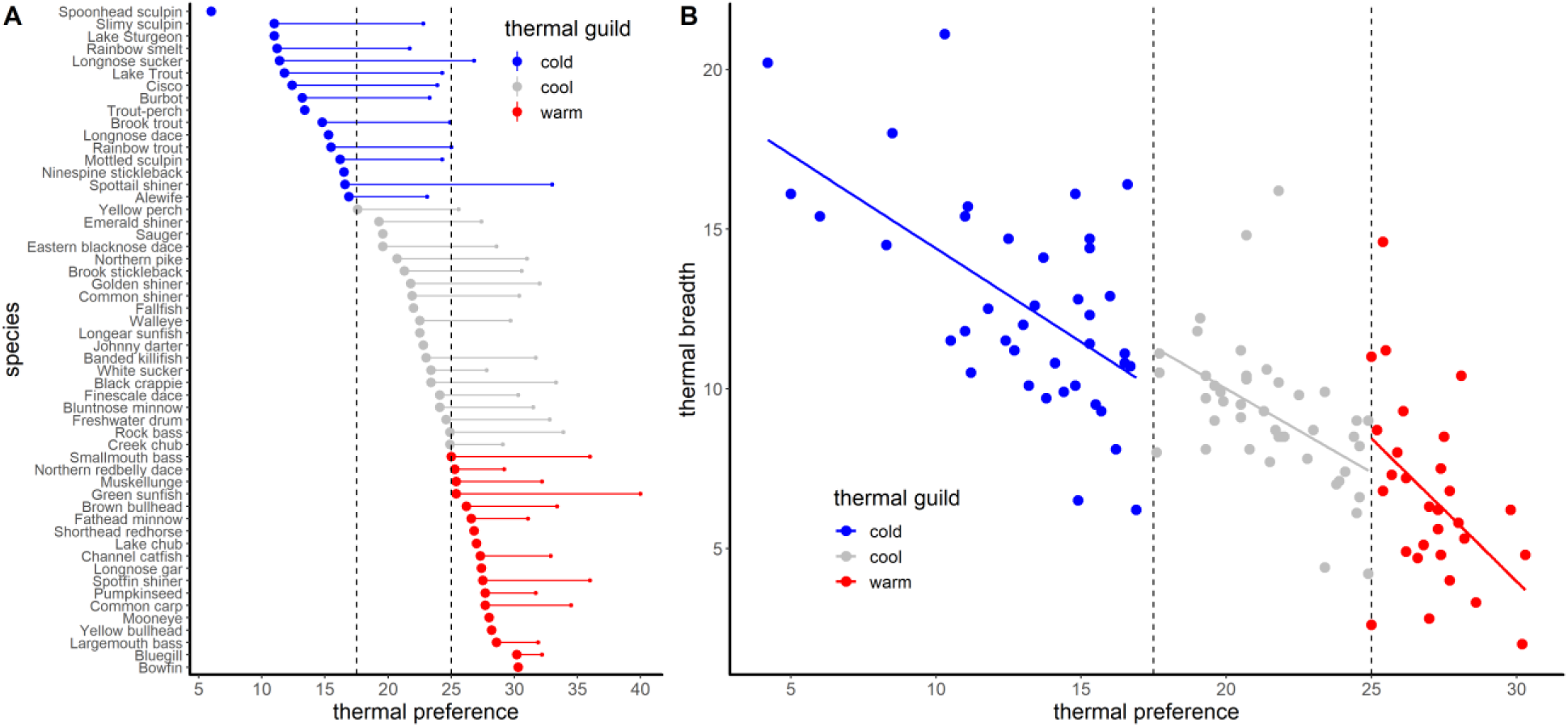
Thermal preferences and thermal niche breadth for Canadian freshwater fishes belonging to three thermal guilds: cold-water, cool-water, and warm-water. A) The continuous distribution of thermal preference (large left point) and upper thermal limit (small right point) for 54 of the most common Canadian freshwater fishes. B) The relationship between thermal preference and thermal breadth (upper thermal limit – thermal preference) for 114 species across the three thermal guilds is negative. These data support thermal performance trade-offs in freshwater fishes. Increased performance at cold temperatures should trade-off with reduced performance at warm temperatures (A) and increased performance at warm temperatures should trade-off with reduced thermal breadth (B). Thermal preference is defined as the temperature that the fish gravitate towards when provided with a broad range of temperatures. Upper thermal limit is the temperature at which 50% mortality occurs in a population. Data for both panels are from Hasnain et al (2018).

Despite winter being a potentially stressful time for all fish, physiological adaptations allow for sustained winter activity in some species. These adaptations include hypertrophy of the heart and liver, increased mitochondrial density and function, improved vision, and high growth efficiency (Shuter et al. 2012; Tschantz et al. 2002). Adaptations to activity in the cold do come with a cost (e.g., greater mitochondrial density can increase standard metabolic rate; Pörtner 2002), but these metabolic costs should be lower for species with lower thermal preferences compared to species with warmer thermal preferences (Shuter et al. 2012). This pattern is suggested by Fig. 2B, showing that thermal breath (i.e. upper thermal limit – thermal preference) narrows as thermal preference increases. The cost of optimizing to a particular temperature is also thought to generate thermal performance trade-offs, meaning a species cannot perform optimally across all temperatures (Kingsolver 2009; Portner 2007). The thermal metrics for North American freshwater fishes provide strong empirical support for such thermal performance trade-offs. First, species with colder thermal preferences have lower upper thermal limits (Fig. 2A). In fact, upper lethal temperatures for many cold-water fish are actually less than the preferred temperatures for many warm-water fish (Fig. 2A). Increased performance at lower temperatures therefore appears to trade-off with reduced performance at higher temperatures (Fig. 1C, 2A). Additionally, increased performance at higher temperatures appears to trade off with lower thermal breadth, suggesting a trade-off between thermal preference and thermal breadth (Fig. 2B). Cold water fish should therefore be more successful across seasons compared to warm water fish, whose performance is limited during winter. We now present two case studies to explore how different capacities for winter activity and trade-offs between activity in open water vs. ice cover manifest in particular species pairs among different (case study #1) and within the same thermal guild (case study #2).

#### Case Study #1

The first case study presents data from a northern temperate lake in Ontario, Canada for smallmouth bass (*Micropterus dolomieu*) and lake trout (*Salvelinus namaycush*), which belong to the warm-water and cold-water thermal guilds, respectively (Fig 2A). This lake lacks an offshore forage fish, meaning the only prey fish for consumption by piscivores are found in the littoral zone. In lakes supporting this type of food web, lake trout and smallmouth bass have overlapping diets that include invertebrates and littoral forage fish (Box 1). Acoustic telemetry data (explained in detail in SI S1) confirm that during cold months, lake trout continue to be active and move inshore, overlapping in the littoral habitat used by smallmouth bass (Fig. 3A, B, C).

###### Box 1. Winter mediates biotic interactions: an empirical case study between thermal guilds

As an example of how winter can mediate the interaction between predator species with differing thermal preferences in a way that could promote their coexistence, we looked at smallmouth bass (warm-water guild) and lake trout (cold-water guild) in Lake of the Two Rivers, Ontario, Canada (45° 34’ 42.6” N, 78°29’ 0.4” W, 274 ha, 38 m maximum depth). New acoustic telemetry data were available for both species that were tracked simultaneously for 1.5 years. Both species are generalist predators that consume both invertebrates and fish. Although lake trout are restricted to cold, offshore waters during summer and smallmouth bass occupy warm, inshore zones, lake trout make inshore forays to consume littoral invertebrates and fish (Guzzo et al. 2017; Tunney et al. 2012). Inshore foraging is especially important for lake trout production in small lakes that lack an offshore forage fish, like our study lake (i.e., where lake trout must move to littoral zones to feed on fish; Vander Zanden and Rasmussen 1996). When present, invasive smallmouth bass have been shown to reduce lake trout reliance on littoral prey compared to lakes that lack smallmouth bass (Vander Zanden et al. 1999). Therefore, lake trout likely experience negative competitive effects from smallmouth bass presence in our study lake (either exploitative or interference).

The acoustic telemetry data were used to calculate both spatial positioning of each species (akin to spatial habitat partitioning through time) and mean activity rates (akin to temporal niche partitioning; see SI S1 for more detail). The spatial data demonstrated clear spatial separation of the two species in summer. Smallmouth bass occupied the warmest, most inshore and shallow waters of the lake and lake trout occupied offshore, cooler water below the thermocline (Fig. 3A, B, C). These tendencies are consistent with temperature-driven, resource partitioning and support its potential role as an important coexistence mechanism for these species (Tonn and Magnuson 1987). As the lake turns over and surface waters fall below 15°C in the fall, however, lake trout ascend in the water column towards the surface and move into inshore, littoral habitats (Fig. 3B, C). During winter, lake trout remain inshore and occupy shallower water than smallmouth bass throughout the winter and early spring. Smallmouth bass moved into slightly deeper water that is warmer than immediately below the ice-water interface (Fig. 3A, B).

Despite the increase in spatial overlap between these two species during fall turnover and winter ice cover, they appear to differentiate in time. The activity rate data provided the important insight that, as the two species converge in space, smallmouth bass have substantially reduced activity levels (Fig. 3D). Mean daily activity rates declined from ∼6-8 m min^−1^ during summer to near-0 during winter for smallmouth bass, but remain comparable during summer and winter for lake trout (Fig. 3D). Activity spikes by lake trout in the fall were likely associated with fall spawning, but otherwise, lake trout remained as active in winter as they were in the spring and summer. Previous findings also demonstrate that warm water fish reduce their activity and foraging under ice (although some swimming and foraging is still possible; Suski et al. 2007), and that cold water salmonids can sustain similar activity rates between summer and winter (Blanchfield et al. 2009).

Seasonal patterns in activity appear to translate to inter-annual patterns in individual growth for these two species. Although growth data were not available for both lake trout and smallmouth bass in our study lake, back-calculated individual growth data from Lake Opeongo (a larger, nearby lake that contains an offshore forage fish) suggest that lake trout are better able to maintain their biomass during years with long winters compared to smallmouth bass (Fig. 4A, B). Although lake trout growth rates vary widely from lake to lake (e.g., depending on the presence or absence of an offshore forage fish; Fig. 4C), lake trout are expected to be more active and capable of feeding than smallmouth bass during winter based on bioenergetic considerations (see SI S3). More data are needed to establish the context dependencies of how winter duration influences relative growth rates between competing species across lakes with different characteristics. However, years with longer winters would be expected to reduce the time for resource acquisition and investment into growth and production of offspring for the warm-water smallmouth bass to a greater degree than cold-water lake trout (Christie & Regier 1988; Giacomini & Shuter 2013; King et al. 1999).

**Figure 3.**
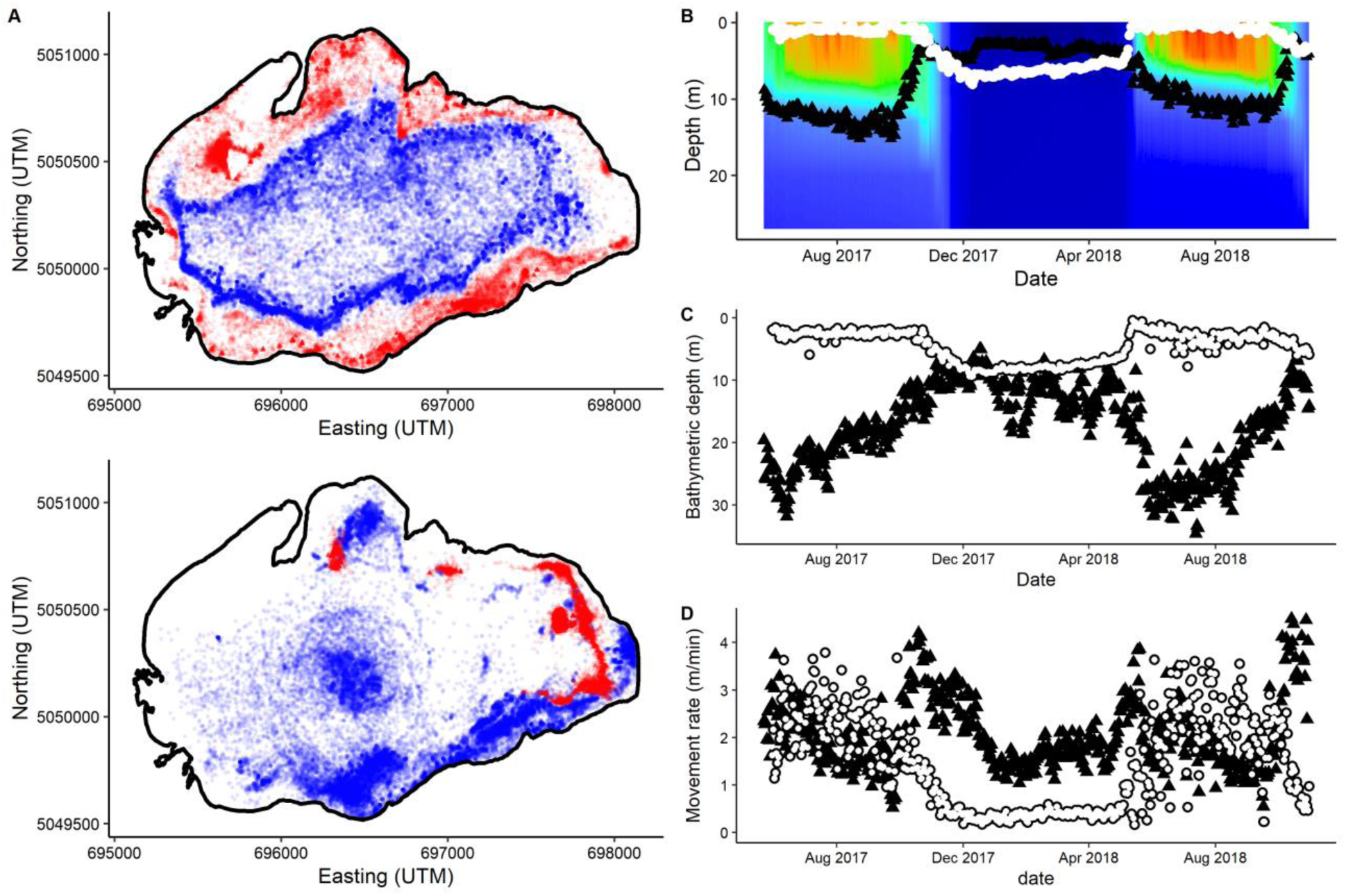
Acoustic telemetry data for lake trout (blue relocations and black triangles) and smallmouth bass (red relocations and white circles) from Lake of the Two Rivers (Ontario, Canada). A) Spatial relocations in summer (top plot; July and August) and winter (bottom plot; January and February). B) Mean daily depth (m) of fish within the water column overlaid over the thermal profile of the lake across the time series. C) Mean daily bathymetric depth (m) (i.e., depth of water over which fish was positioned). D) Mean daily activity rates (m min^−1^). For details on the acoustic telemetry data and analyses, see SI S1.

However, smallmouth bass, but not lake trout, reduce their activity rates in winter as spatial overlap increases (Fig. 3D), suggesting temporal niche partitioning occurs between these two potential competitors (Box 1). Differences in growth rates further suggests that different seasons favor different species (Fig. 4, Box 1), consistent with the necessary conditions for temporal fluctuation-depending coexistence.

**Figure 4.**
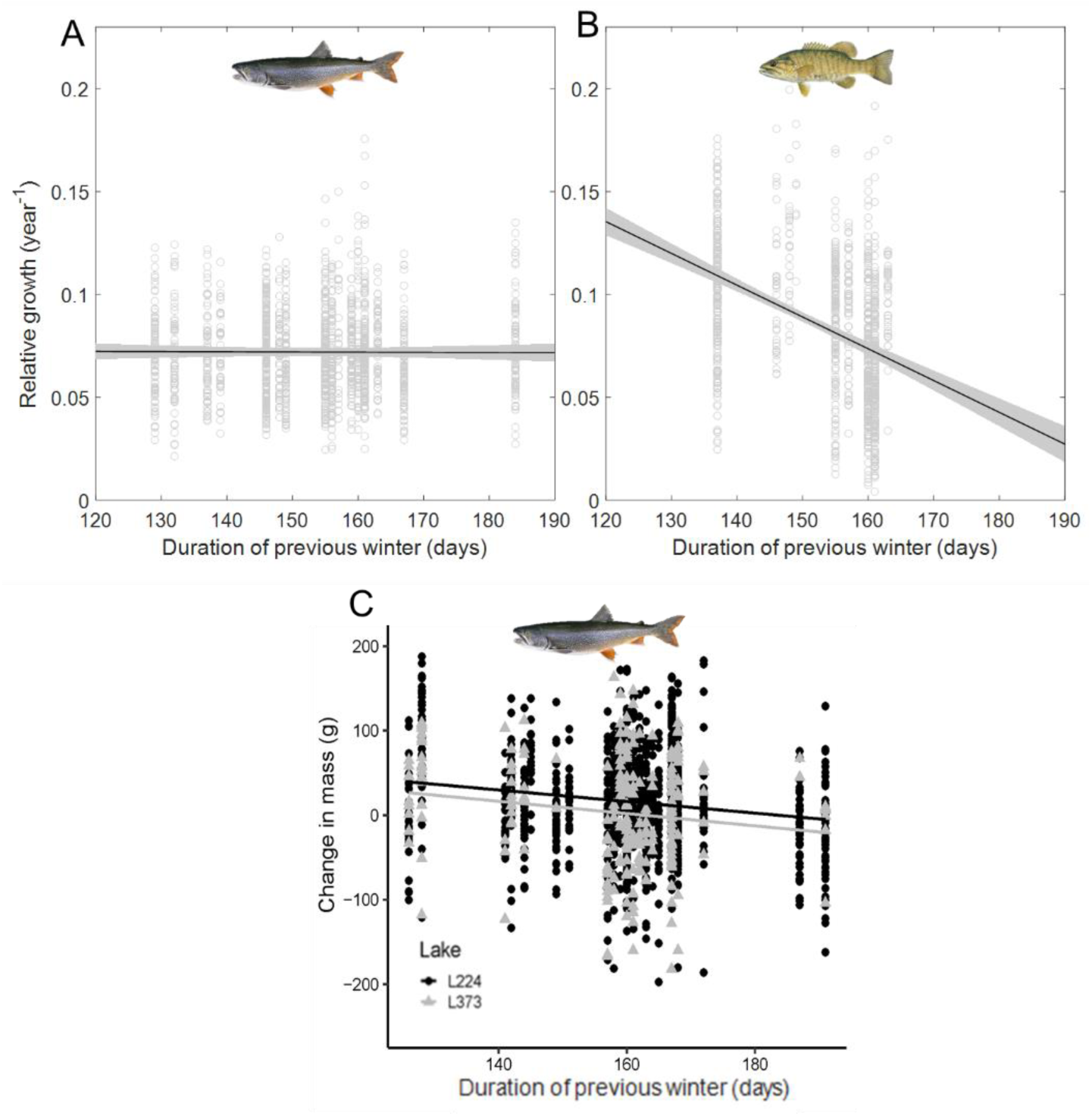
Individual growth of lake trout (A, C) and smallmouth bass (B) in response to inter-annual changes in winter duration (days). Data from Lake Opeongo, Ontario, Canada (A, B; cm per year relative to maximum length) demonstrate that lake trout are more tolerant to winter duration than smallmouth bass. In smaller lakes lacking an offshore forage fish (Experimental Lakes Area Lake 224 and Lake 373), like our study lake described in Case Study #1 (Box 1), however, individual lake trout mass (g) declines in years with the longest winters (C). See SI S2 for details on growth calculation and data analysis.

#### Case study #2

The second case study presents data from a northern temperate lake in Northwest Territories, Canada for lake trout and burbot (*Lota lota*), which both have cold thermal preferences (Box 2). These species segregate spatially throughout much of the year (Fig. 5A, B) but have similar diets both being piscivores (Guzzo et al. 2016b). In accordance with their colder thermal preferences, previous work has illustrated that both of these species can be active during winter (Blanchfield et al. 2009; Hölker et al. 2004), suggesting competition could be high throughout the year, including under winter ice cover. Indeed, acoustic telemetry data demonstrate that these two species overlap in their patterns of annual activity (i.e., both reduce activity in summer and maintain activity in winter, Box 2, Fig. 5C). Regardless, lake trout and burbot clearly show temporally distinct niches that likely reduce competition. Lake trout spawn in the fall and are known to forage actively in the spring, whereas burbot are actively foraging and spawn in the winter (Hölker et al. 2004, Guzzo et al. 2016b). Shorter (darker) days and increased ice and snow cover of late winter might, therefore, favor the typically nocturnal burbot over the more visually-reliant lake trout (Box 2). As the ice thins and melts in the early spring, lake trout and burbot would be expected to exploit suitable conditions for foraging and growth, but before warm-adapted species become active (e.g., smallmouth bass; Box 1).

###### Box 2: Empirical case study of winter habitat use and activity rates within a thermal guild

To evaluate how species that share thermal preferences differentiate in winter, we examined lake trout and burbot using available acoustic telemetry data from Alexie Lake (62°40′36.59″ N, 114° 4′22.76″W), located approximately 30 km north east of Yellowknife, Northwest Territories (NT), Canada. Alexie Lake is a medium-sized (402 ha, maximum depth 32 m) oligotrophic lake that thermally stratifies in the summer and experiences approximately 6 months of ice cover annually. Lake trout and burbot both belong to the cold water thermal guild and are piscivores, but separate spatially in their habitat use (Guzzo et al. 2016b). Acoustic telemetry data (see SI S1 for detail) indicate that lake trout are pelagic, spending the majority of their time around 10 m depth, while burbot are benthic and spend time on the bottom in shallower water (Fig. 5A, B). Seasonal patterns in mean daily activity rates broadly overlap because both species exhibit their minimum activity rates in the summer (Fig. 5C), when they both occupy deeper water (Fig. 5B). However, the timing of maximum activity rates clearly diverge, reflecting differences in their life history. Lake trout peak in their activity in the fall while burbot peak in the winter (Fig. 5C). While lake trout do remain active in winter, because winter activity rates stay above minimum values observed in summer, activity rates slowly decline as the winter progresses in this sub-Arctic lake (from December to March; Fig. 5C). This is, however, not the case for burbot, which spawn in the winter under the ice (Cott et al. 2013). Burbot remain as active in late winter as early winter, and actually increase their activity through winter as their spawn time approaches (Fig. 5C).

Lake trout and burbot are additionally known to diverge in the daily timing of their foraging. Lake trout feed most during the day and burbot during the night (Guzzo et al. 2016b, Cott et al. 2015). While this would prevent interference competition from occurring, exploitative competition is still possible if one species is suppressing the resources available to the other species. Lake trout are known to be visual predators (Blanchfield et al. 2009; Vogel & Beauchamp 1999). Alternatively, the photophobic behavior of burbot, which successfully and exclusively forages in the dark (Cott et al. 2015), could suggest they are more effective predators and competitors during later parts of the winter, when light would be more limited by both reduced photoperiod and more snow and ice build-up. We do not have prey capture rates or winter diet information to explore this idea further. However, a greater foraging advantage provided by dark winters to burbot could explain their sustained activity rates during this part of the season compared to lake trout. Trade-offs between successful foraging in bright vs. dark parts of the year could therefore be operating to favor different species at different times (Helland et al. 2011; Vogel & Beauchamp 1999), similar to the thermally driven trade-off that seems to favor warm-adapted species in summer and cold-adapted species in winter (Box 1). Here, more visual species could be favored by brighter seasons (fall, early winter and spring) relative to species that forage better in dark, late winter conditions. Again, each species having their time of greatest success is the key to fluctuation-dependent coexistence. This example suggests that species can partition both among but also within a given season (i.e., early vs. late winter).

**Figure 5.**
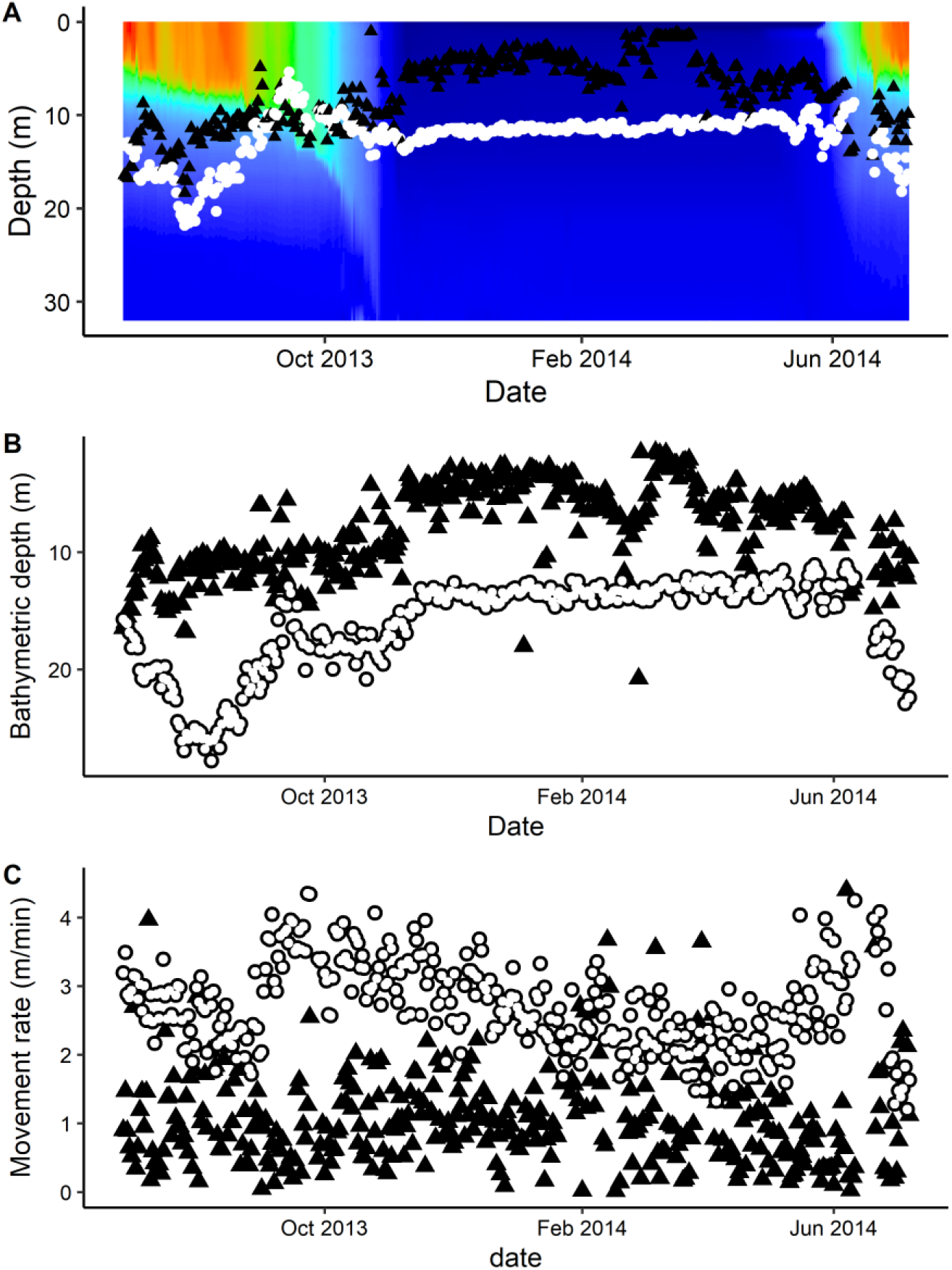
Acoustic telemetry data for lake trout (white circles) and burbot (black triangles) from Alexie Lake (Northwest Territories, Canada). A) Mean daily lake depth (m) of fish within the water column overlaid over the thermal profile of the lake across the time series. B) Mean daily bathymetric depth (m) (i.e., depth of water over which fish was positioned). C) Mean daily activity rates (m min^−1^). See SI S1 for more information.

#### Other examples of seasonally variable biotic interactions

Fluctuating temperatures can alter the identity of the competitive dominant between other pairs of fish species, too. For example, European perch (*Perca fluviatilis*) had higher prey capture success and were competitively superior at cooler temperatures closer to their optimum, compared to roach (*Rutilus rutilus;* Persson 1987). Conversely, roach performed better at warmer temperatures closer to their optimum (Persson 1987). Other examples from the literature confirm that seasonal cycles of photoperiod and light availability can facilitate cycles of competitive advantage among fishes with similar thermal preferences but divergent requirements for light. In European lakes, brown trout (*Salmo trutta*) and Arctic charr (*Salvelinus alpinus*) are both coldwater salmonids that overlap in their littoral habitat and dietary preferences (Amundsen & Knudsen 2009). Brown trout are considered the superior competitor and appear capable of restricting Arctic charr from accessing littoral habitats during summer (Eloranta et al. 2013). Under ice cover, however, Arctic charr move into the littoral habitat (Amundsen & Knudsen 2009) and appear to be the superior competitor, having a higher capacity for successful foraging in the dark compared to brown trout (Helland et al. 2011).

The examples we have highlighted so far illustrate that winter behavior can vary both among and within thermal guilds. Organisms cannot succeed and perform best under all conditions, especially when conditions vary across seasons (Kingsolver 2009, Pörtner 2002). For example, species that invest heavily in rapid growth do not tend to invest in physiological adaptations for coping with sub-optimal periods (Angert et al. 2009; Chesson & Huntly 1997). Performance at high temperatures can trade-off with performance at cold temperatures, and having a wide thermal breadth can trade-off with maximum performance at optimal temperatures (Kingsolver 2009). Physiological adaptations that favor active winter foraging and a wider thermal breadth could, therefore, limit these species ability to use the warmest, brightest, and more productive region of lake (i.e., nearshore littoral zone) during summer (e.g., Fig. 2). Next, we discuss how these trade-offs could play a role in coexistence based on modern theory.

#### Coexistence in time and the role of winter

Trade-offs that favor different species at different times and generate differential demographic responses to the environment are the basis for fluctuation-dependent coexistence (Angert et al. 2009; Chesson 2000; Miller and Klausmeier 2017). We lack data on the competitive response, functional response, or average fitness of our case study species to properly test for a particular coexistence mechanism in northern temperate lakes. However, our empirical examples (see Boxes 1 and 2) appear to meet the three ingredients of the storage effect. Fish are expected to demonstrate differential demographic responses to the environment (i.e., condition #1) because their activity, growth and reproduction are favored and cued by different environmental conditions (Shuter et al. 2012). Winter-active species might experience weaker environment-competition covariance (i.e., condition #2) if competition weakens during winter, when otherwise superior competitors become inactive or less successful foragers and competitors. Longer life expectancies would buffer fish population growth from ‘bad’ times when the environment is not favorable and competition is high (i.e., condition #3). Of course, other mechanisms are possible, and the storage effect and relative nonlinearity are not mutually exclusive. For example, different growth responses to winter duration could also reflect relative nonlinearity of species functional responses. A trade-off between growth capacity in open water and winter activity could also suggest winter acts as an equalizing mechanism by reducing the population growth of the species with higher growth capacity.

Based on the above arguments, our perspective is that winter, which is historically underrepresented in ecology, could play an important role in weakening biotic interactions (like competition) in ways that promote coexistence through a variety of theoretically supported mechanisms. We are not arguing that winter is the only season that is important for coexistence; competition might actually peak during other seasons when spatial overlap is highest (e.g., during fall and spring when lake trout and smallmouth bass cross paths, Fig. 3B, C). Seasons other than winter can also be harsh and stressful to fish (e.g., summer can cause thermal or oxidative stress), and fish clearly diverge in their foraging behavior and life history in lakes that lack ice-covered winters, meaning that winter is not a prerequisite for fish to coexist. Our overarching goal, here, is to bring attention to two key points:

1. Biotic interactions between species can change dramatically throughout the year with seasonal changes in environmental conditions.
2. Ice-covered periods, which are broadly perceived as a stressor for all fishes, could also play a unique role in generating differential responses among competing species.

While all fishes are expected to be active during open water periods and generally able to find and use habitat within optimal temperatures, the unique conditions of winter in ice-covered lakes seem to result in divergent behaviors that include both winter active and inactive species. The persistent cold, dark, low-resource, or low-oxygen conditions of winter drive species with warmer thermal preferences to suppress their activity rates and reduce their foraging (by accumulating lipid reserves before winter or relying on energy conservation due to suppressed metabolic rates or both; Mackereth et al. 1999; Secor & Carey 2011, Shuter et al. 2012). Some evidence suggests, for example, that warm-water fishes in southern temperate lakes with warmer, shorter and more sporadic winters do not enter a period of consistently reduced activity (Fullerton et al. 2000). Dark periods that reduce the performance of species tailored to thrive in bright, open waters (Helland et al. 2011; Vogel & Beauchamp 1999) hinges on stable, lasting snow and ice cover that suppress light beyond that which is associated with photoperiodic reductions. If winter structures biotic interactions in ways that weaken competition, increased water temperature and light (i.e., decreased snow and ice) due to climate change could alter competitive advantages by increasing activity rates or prey capture success for species that would otherwise have suppressed winter performance. This is akin to species increasing their niche overlap in time. Given its predictability to organisms, winter could be a strong generator of divergent ecological behaviors among species (i.e., timing of reproduction, growth, activity) and performance trade-offs. If such trade-offs underpin coexistence and tend to organize around predictable variation in space (e.g., littoral versus pelagic) and time (e.g., winter versus summer), it is critical that we deepen our understanding of ecological processes occurring during winter. We now consider how winter and temporal niche divergence might play a widespread role in coexistence even in systems where species also spatially partition resources.

### A case for multiple coexistence mechanisms

To this point, we have focused on coexistence that arises by species partitioning along seasonal axes by diverging in the timing of their activity. However, biodiversity in complex, natural systems is almost certainly sustained by species partitioning along multiple niche axes. Mobile species are readily able to avoid competition by spatiotemporally partitioning their environment (e.g. by moving to particular locations during certain seasons that contain lower densities of competitors; Jeltsch et al. 2013). Given the high mobility and generalist diet of many fishes, spatial and temporal niche partitioning likely operate in tandem. For example, mobile aquatic taxa shift their habitat seasonally in response to changes in temperature and prey density (Diez et al. 2018; Holbrook and Schmitt 1989) or select for prey types not shared by their competitors during each season (Amundsen & Knudsen 2009; Hayden et al. 2015). Even in our case studies, competing species pairs diverge in their annual patterns of activity but also their habitat use (Box 1, 2). Coexistence in fish communities could therefore be maintained by divergence in both time and space.

Recent theoretical and empirical studies have also concluded that species diverge along multiple niche axes, and that multiple coexistence mechanisms likely operate simultaneously to sustain nature’s diversity (Chesson et al. 2012; Chesson 2018, 2019; Ellner et al. 2016; Zepeda & Martorell 2019). Using different resources and using the same resource at different times would both be expected to stabilize coexistence for the same reason: both dampen the strength of inter-specific relative to intra-specific competition (Chesson 2000; Chesson & Huntely 1997; Chesson 1985). The existence of multiple coexistence mechanisms means that, even if species exhibit some overlap in temporal activity patterns, species can still coexist by partitioning resources in space at any given time. Performance trade-offs (e.g., between summer growth capacity and winter activity) need therefore not be perfectly balanced to play a role in coexistence. For example, species that are well-adapted to warm productive periods could also be capable of low-level winter activity, which would drive some overlap in annual activity patterns with a winter-active species. Cold-water species that are capable of winter activity could have similar activity rates to warm-water species in summer, albeit in different habitats (e.g., lake trout and smallmouth bass, Box 2). Divergence along multiple niche axes could make coexistence possible in these situations, where some overlap occurs along one axis.

Importantly, if seasonal variation and harsh winter periods drive species to diverge along any niche axis, whether that be in the timing of foraging, reproduction and growth, or in their habitat and resource selection, it could be an under-recognized player operating in a suite of coexistence mechanisms. In other words, winter might play a role not only in driving divergence in activity patterns, but also in species selecting for different resources and habitats (Schoener 1982). Shorter, weaker winters already arising in northern temperate lakes (Weyhenmeyer et al. 2011, Sharma et al. 2019) could therefore threaten coexistence if they cause species to converge in their niches, either by causing species to become more winter-active when they would otherwise be inactive, or by causing species to overlap in their resource use. As a step towards considering the biodiversity consequences of ice cover loss on northern temperate lakes, we developed new theory that explores under what contexts winter might play the most important role in coexistence.

### Winter’s context dependency: when and where does winter matter most for coexistence?

Different winter behaviors have different consequences for individual survival, growth, and therefore, population dynamics (Hurst 2007). Foraging in the winter increases both metabolic costs and potential energy gain, which could have either positive or negative consequences for growth (Byström et al. 2006; French et al. 2014; Amundsen & Knudsen 2009; Cunjak et al. 1987). In lake trout, for example, long winters appear to have minimal impact on individual growth in lakes that contain offshore forage fish, yet suppress lake trout growth in lakes lacking offshore forage fish (Fig. 4A, C). Access to a high-quality prey during open water months, which increases lake trout growth and condition (Cruz-Font et al. 2019), could carry over and help promote more sustained growth in winter, although more work is needed to explore this idea. Compared to winter-active species, winter-inactive species suppress metabolic costs through reduced activity and feeding and therefore have little capacity for overwinter growth (Micucci et al. 2003). Food availability, predation, disease, starvation, and competition could all drive variation in overwinter growth and mortality within and among winter active and inactive strategies (Garvey et al. 2004; Hurst 2007). In small lakes, for example, resources may be limited and competition more amplified due to higher spatial overlap between competing species (Hayden et al. 2014, McCann et al. 2005). The consequences imposed by winter should therefore be expected to vary substantially among different fish species (Shuter et al. 2012), and from lake to lake among different populations.

Given the paucity of winter data, it is unknown if and how fish growth and competition during winter vary with characteristics like lake size and productivity in ways that influence coexistence. To begin to address this, we developed new theory that explores how coexistence in a seasonal environment is dependent on the duration of winter under cases of both weak and strong competition (as might be expected in large and small lakes, for example). Given the variable outcomes of winter for fish activity and growth across species and populations, we also consider coexistence outcomes when the two competing species have either synchronized (i.e., both decline in winter to various degrees) or asynchronized growth dynamics (i.e., the winter-adapted species grows in winter, Fig. S2). Below, we highlight two important points from this theory, which is explained in Box 3.

##### Box 3. Theoretical consideration of how winter influences coexistence

We used the classical 2 species Lotka-Volterra competition models, but explicitly consider dynamics in two discrete seasons, open water and winter. Open water (*f_S_*) and winter (*f_W_*) are modelled as fractions summing to 1 (*f_S_ + f_W_ =1*) and each species is modelled with season-specific parameter combinations. In this way models are integrated over numerous years but within each year they sequentially follow first open water then winter parameters corresponding to the given seasonal fractions within the year. The seasonal fractions allow us to change the proportion of the year that is winter (e.g., reduce *f_W_* to mimic warming). The open water and winter models for species X_1_ and X_2_ are as follows:

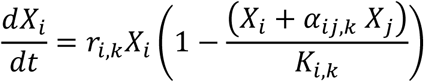

where *i* and *j* are the competing species, *r_i,k_* is the intrinsic rate of population growth for species *i* in season *k* (i.e., open water or winter), *K_i,k_* is the carrying capacity (a surrogate for habitat productivity) for species *i* in season *k*, and *α_ij,k_*, the competition coefficient, is the competitive effect of species *j* on *i* in season *k* (see Table 1 for parameters). Note that many competition models nondimensionalize the parameters such that interspecific competition is scaled to *α_ij,k_/K_i_* (Chesson 2000). In our model, we wanted to alter carrying capacity explicitly (*K_i_*) in order to match our empirical observations that the accessibility of resources in a given habitat changes seasonally (Guzzo et al. 2017). In our case, competition can change indirectly as a result of *K_i_* (lower *K_i_* reduces resource availability and effectively increases inter- and intraspecific competition) or directly by changing the *α_ij_* (Table 1).

With respect to the seasonal environment, we make the following two assumptions:

1. Winter is less productive than open water (i.e., *K_i,S_* > *K_i,W_*)
2. Maximal growth rates are smaller in winter than open water (i.e., *r_i,S_* > *r_i,W_*)

For the two species, following the patterns discussed in the main text, we assume the following trade-offs in the parameters:

1. Species 1 has higher growth rates in open water (*r_1,S_ > r_2,S_*) and lower growth rates in winter (*r_2,W_ > r_1,W_*), compared to species 2.
2. Species 2 growth rate remains more constant throughout the year.
3. Species 2 is a stronger competitor in winter (*α_12,W_ > α_21,W_*), but Species 1 is a stronger competitor in open water (*α_21,S_ > α_12,S_*).

With the above assumptions, we then consider four different competition cases where we alter the strength of interspecific competition to explore how winter length (*f_W_*) influences coexistence of the two species under different, empirically motivated scenarios of strong vs. weak competition in each season (Table 1). We explored the qualitative outcomes for coexistence under both synchronized and asynchronized species dynamics (see SI S4).

*Case 1. Competition is relatively weak in both seasons (Weak-weak, Fig. 6A).* Here, weak-weak refers to competition coefficients that yield coexistence conditions in both open water and winter. Weak competition could arise in productive or large lakes where carrying capacity is high and/or species spatially separate in their habitat and resource use, or when species diverge in their winter activity rates (one is active and one is inactive).

*Case 2. Competition strengthens in summer but remains weak in winter (Strong-weak, Fig. 6B).* Here, strong-weak refers to competition coefficients that yield competitive exclusion conditions in summer and coexistence in winter. Competition could strengthen (relative to case 1) in summer in lower productivity or smaller lakes. Winter competition could remain weak for the reasons listed above in case 1 (high carrying capacity or high niche partitioning).

*Case 3. Competition remains weak in summer but strengthens in winter (Weak-strong, Fig. 6C).* Weak-strong refers to competition coefficients that yield coexistence conditions in summer and exclusion in winter. Here, species have a high carrying capacity or partition resources in summer that weakens competition, as in case 1 above. During winter, however, either carrying capacity declines or species niche overlap increases, amplifying competition compared to case 1. We might expect reduced winter carrying capacity in lower productivity lakes and higher niche overlap in smaller lakes or in cases where both species are actively foraging during winter.

*Case 4. Strong competition in both summer and winter (Strong-strong, Fig. 6D).* Strong-strong refers to competition coefficients that yield exclusion conditions in both summer and winter. In smaller lakes with less ice cover, we imagine that competition could be strengthened relative to case 1 in both summer (due for example to increased resource overlap) and winter (due for example to greater synchrony in annual activity patterns).

#### Winter duration and the strength of competition impact species coexistence

The length of winter impacts species coexistence generally in that it can drive the loss of diversity or, at the least, produce large changes in the density of competing species based on our theory (Fig. 6). Our theory also suggests the intriguing case where seasonality can be entirely responsible for coexistence given the empirically supported trade-offs whereby the species that is most successful in winter is less successful in open water (Fig. 2; Portner 2007; Lancaster et al. 2017). Note, the strong-strong case by definition does not give coexistence without seasonality (Fig. 6D). This **winter-mediated species coexistence** is interesting because it suggests climate change that alters winter length and ice cover duration, as predicted across northern temperate lakes (Weyhenmeyer et al. 2011), even modestly, ought to potentially significantly and rapidly drive the loss of winter-adapted species (Fig. 6C). This same sensitivity to seasonal length also occurs for other cases (Fig. 6), so the impacts of season are potentially potent even where only one season mediates coexistence.

**Figure 6.**
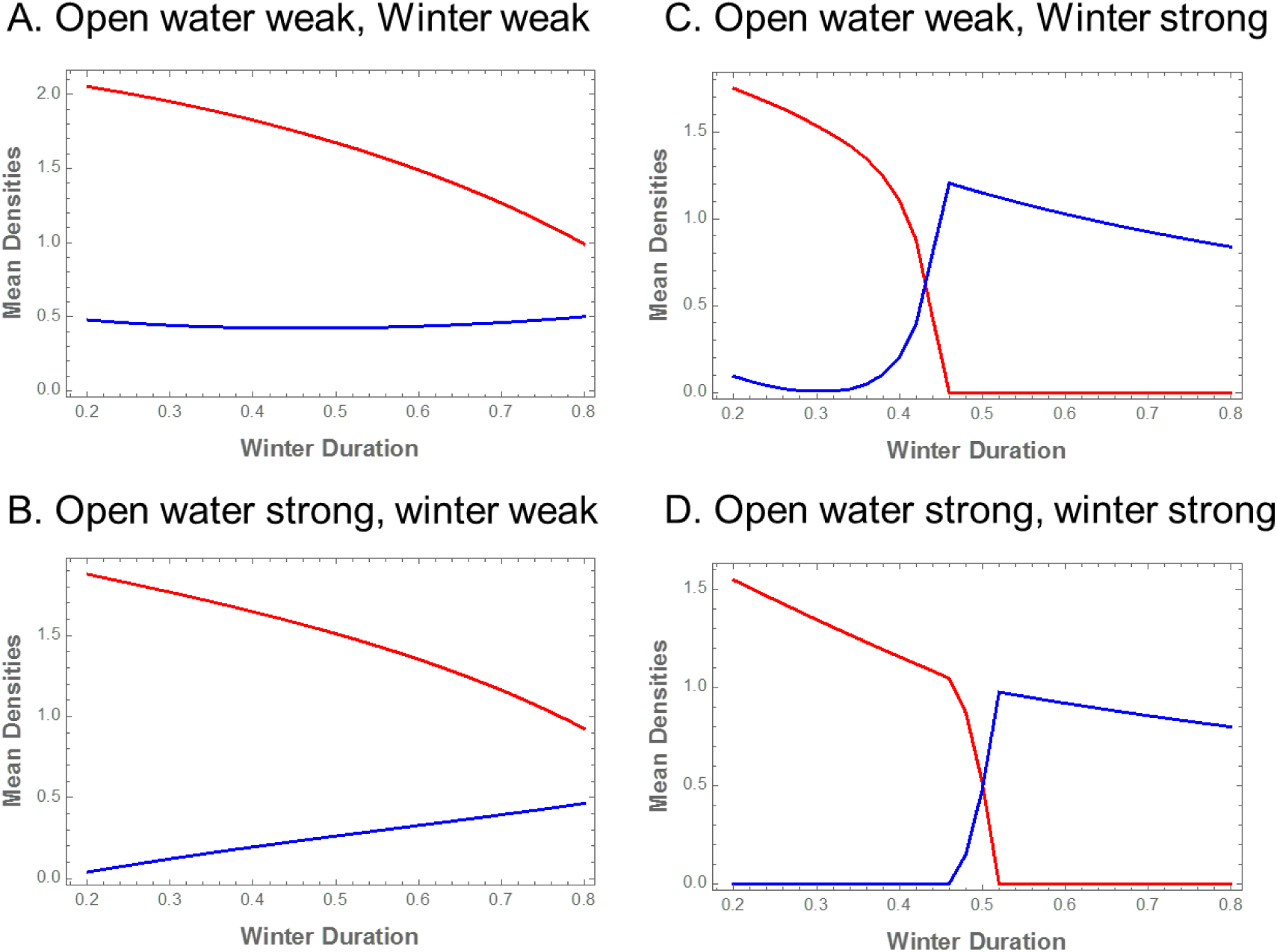
Densities of two competing species, one adapted to open water (red) and one to winter (blue) across a gradient of changing winter duration. Four cases were considered, where competition is either weak or strong in each season. Weak refers to a case where the two species coexist in a given season and strong refers to competitive exclusion in a given season (see Box 3 for detail).

Coexistence between two species was, however, sensitive to the strength of competition. When competition was weak in both seasons, the two species coexisted across a broader range of winter durations (Fig. 6A). Weak competition might be expected in large lakes, where spatial overlap among competitors could be low, and where species have opportunities to more effectively partition resources (Hayden et al. 2014, McCann et al. 2005). For example, lake trout reduce their reliance on littoral foraging as lake size increases (Tunney et al. 2012), which could reduce their resource overlap and potential competition with littoral fishes like smallmouth bass (Vander Zanden and Rasmussen 1996, Vander Zanden et al. 1999). Large lakes might also be expected to have more habitat refuges for cold-adapted fishes, because larger lakes are typically deeper. As winters warm, diet shifts to alternate prey or different habitats in order to avoid competition could therefore be more probable in large lakes. Such systems, where adaptive capacity is high, could be less sensitive to biodiversity loss under warmer winters.

Cases in which competition during winter was strong narrowed the region of coexistence (Fig. 6C, D). In these strong cases, the winter-adapted species still had a growth and competitive advantage during winter (i.e. higher *r* and α12,W; Table 1). Reducing the duration of winter easily drives the winter-adapted species to extinction in such cases when winter competition is strong, such that winter-adapted species depend on this time period for maintaining coexistence (Fig. 6C, D). In a simplified sense, the theory therefore suggests that the role of time and space in mediating coexistence may be context-dependent. More specifically, when an ecosystem is spatially constrained with heightened interactions within a trophic level (e.g., small lakes *sensu* McCann et al. 2005), one may reasonably expect that temporal trade-offs play an increasingly large role in mediating coexistence. Given this, we may predict that winter (and niche divergence through time) could play an increasingly large role in spatially constrained ecosystems for competing mobile species (e.g., top predators in small lakes and islands that overlap significantly in space). If so, the global warming implications may be dire for predator diversity in small temperate lakes. Future work is therefore warranted to begin to develop new competition theory to understand when temporal vs. spatial coexistence mechanisms may be expected to operate, even within the same ecosystem, and including mobile species (e.g., see Chesson 1986 for an example relevant to species with dispersing larva and sedentary adults).

**Table 1.** Parameter values used to (1) generate trade-offs between a warm adapted and cold-adapted species, and (2) to create strong and weak competition scenarios during two seasons.

| Summer |  |  |  | Winter |  |  |  |
| --- | --- | --- | --- | --- | --- | --- | --- |
| Strong | $a_{12} = 0.5$ | Weak | $a_{12} = 0.5$ | Strong | $a_{12} = 1.3$ | Weak | $a_{12} = 1.1$ |
| | $a_{21} = 0.8$ | | $a_{21} = 0.8$ | | $a_{21} = 1.1$ | | $a_{21} = 0.3$ |
| | $r_1 = 1.7$ | | $r_1 = 1.7$ | | $r_1 = 0.3$ | | $r_1 = 0.3$ |
| | $r_2 = 1.2$ | | $r_2 = 1.2$ | | $r_2 = 1.0$ | | $r_2 = 1.0$ |
| | $K_1 = 2.0$ | | $K_1 = 2.5$ | | $K_1 = 0.25$ | | $K_1 = 0.9$ |
| | $K_2 = 1.5$ | | $K_2 = 2.4$ | | $K_2 = 0.7$ | | $K_2 = 0.7$ |

#### Winter can be bad or good for fish growth and still impact coexistence

The qualitative outcomes of the four cases were not dependent on whether the dynamics were synchronized or asynchronized between the two species (for an example of these dynamics see SI S4, Fig. S5). Coexistence is therefore possible between species exhibiting a trade-off between maximizing open water growth and winter activity, regardless of whether winter is ‘bad’ or ‘good’ for the population growth of the winter-adapted species. As long as the more winter-adapted species is more buffered from negative effects than the summer-adapted species, the two species can coexist. Relative non-linearity (i.e., a fluctuation-dependent coexistence mechanism) also has this feature of certain contexts (e.g., low resource densities) being universally bad for coexisting species (Armstrong and McGehee 1980). Our theory hints that winter need not be a time of positive growth to promote coexistence. On the contrary, winter might still be ‘bad’ for individual and population growth but ‘good’ for biodiversity as long as species respond differently to environmental variation.

### Future work

Northern temperate regions currently have stable temperatures under ice, ensuring predictable metabolic costs for fishes (Garvey et al. 2011) and setting up the conditions for divergent behavioral strategies (i.e., winter active vs. winter inactive; Shuter et al. 2012). The potential for climate change to converge life history and behavioral strategies in ways that intensify competition and threaten biodiversity is receiving increasing attention (Lancaster et al. 2017). Shorter or intermittent ice-cover due to warming could increase the activity of species that otherwise rely on inactivity and energy conservation (e.g., Fullerton et al. 2010), simultaneously reducing the competitive advantage of cold-water species during this period. Species coexistence could further be threatened by increased stochastic environmental variation also expected under a changing climate (Gravel et al. 2011). From an energetic standpoint, warmer, more variable temperatures will be more demanding on ectotherms (Williams et al. 2018). Energy losses over winter have direct effects on the performance and reproduction of fish in the growing season (Hurst 2007). Some evidence has already linked shorter winters with reduced reproductive success of a spring spawning, cool water fish (Farmer et al. 2015). On the other hand, warmer winters might benefit the growth of all fishes if the increase in metabolic demand for resources is met (Brodersen et al. 2011) and, at least temporarily, increase species richness through regional shifts in the species pool (Lancaster et al. 2017). This will not necessarily impinge on coexistence, as long as each species still has its opportunity to succeed under the new conditions of longer growing seasons and warmer temperatures (Lancaster et al. 2017).

Predicting how regional changes in climate conditions will balance with local level shifts in species interactions to ultimately influence species biodiversity is a complex task. Whether northern temperate lakes will eventually lose cold-adapted species and gradually transition into southern temperate lake communities under climate change remains to be seen. We advocate for more directed theoretical and empirical work that considers how species behavior, growth, and reproduction change through seasons and years, including during ice-covered winters, in ways that ultimately influence biotic interactions.

A major challenge of such efforts is measuring competition in the field. However, empirical studies can apply a range of tools for mapping how species interactions change seasonally and from year to year (e.g., across years with short and long winters). Acoustic telemetry to measure the 3D movements and activity of fish in the wild particularly promising (Cruz-Font et al. 2019), especially when coupled with repeated sampling to obtain stomach contents or tissue for dietary analysis (e.g., via stable isotope or fatty acids; Guzzo et al. 2017). Studies have also successfully used netting to capture information about species spatial distributions and overlap through time (Helland et al. 2011). Combined with experimental manipulations of species and resource densities, changes in the strength of competition can be measured between species pairs across changing conditions (including during simulated winter conditions; Hellend et al. 2011).

Winter could also have far-reaching consequences that operate through carry-over effects in other seasons. Species should adaptively anticipate winter’s onset and behave accordingly in the spring, summer, and fall. This includes foraging to ensure sufficient resources are acquired for successful spawning and overwinter survival (Plumb et al. 2014). How successful this foraging is could then influence behaviors and survival during winter. Fall spawning fish that have depleted more of their energy reserves immediately before winter, for example, might be forced to forage more than a spring spawning fish that accumulates surplus energy reserves at this time (Shuter et al. 2012). The ability of a warm-water fish to accumulate energy prior to winter and to locate suitable habitat and conserve energy during winter could dictate its level of winter activity (Bystrom et al. 2006). A thorough consideration of how winter shapes ecological processes should therefore include connections across seasons.

Finally, spatial sampling across different lakes is also necessary to explore the context dependency of winter on ecological processes. Even though sampling the same lake over time may be the best option for quantifying the local effects of winter duration on a system, having a range of lakes sampled through time would allow one to uncover trends in species behavior and interactions across a gradient of lake conditions. We expect lake size and food web structure, including the presence of high- vs. low-quality prey, to be key mediators of how winter influences fish activity, growth, and interactions with competitors. Only by sampling through time across a gradient of lake size, production and food web structure can we begin to understand how local context shapes species responses to winter. Careful attention should be taken in designing such multi-lake studies because in addition to winter duration, a host of other variables can differ among lakes (e.g., productivity, food web structure) that can impact species behavior and interactions.

Although we have focused on species, individuals can also play an important role in coexistence (Clark 2010). Individual fish within a species can have vastly different strategies and behaviors (Mccucci et al. 2003). This individual-level focus remains an important avenue for future work.

We have also stressed the role of competition for coexistence within a trophic guild, due to our focus on top predatory fishes. But seasonal diet switches by these generalist top predators also have important consequences for coexistence within the prey guild (Chesson 2018). For example, flexible diet switches of predators away from declining or inaccessible prey, or suppressed winter activity by a predator, during certain seasons could be important for allowing low density prey to recover (McMeans et al. 2015). Therefore, stabilizing coexistence mechanisms that weaken inter-specific competition within a trophic guild could also dampen otherwise strong predator-prey interactions and stabilize whole food webs. A general framework for how seasonality and harsh winters influence biotic interactions should include individuals, species, and interactions within the community and the larger food web (McMeans et al. 2016; Williams et al. 2018).

A greater focus on seasonality that includes winter periods is warranted. Human activities are broadly homogenizing ecosystems, undermining the heterogeneity that underlies biodiversity. This homogenization is apparent in space (e.g., landscapes transformed for agriculture) and in ecological communities (e.g., the redistribution of species). The climate-driven loss of ice-covered periods on lakes could similarly be viewed as a type of homogenization in time. Weakened winters will make more of the year warmer, brighter, and ice-free, similar to open water periods. If diversity is generated and maintained by species filling niches in time, as the environment fluctuates, homogenization in time could have similar negative effects for biodiversity as homogenization in space (Sabzo et al. 2016). Understanding and predicting the outcomes of altered seasonal conditions demands a more widespread consideration of how temporal variation broadly, and historically understudied winter periods specifically, shape ecological processes that ultimately influence biodiversity maintenance.

#### Predictions

We suggest the following predictions to guide future work:

1. Biotic interactions change through time. Coexisting species diverge in their response to seasonal variation. Species can select for different habitats or resources, or vary in their activity and competitive ability, during different seasons. Differential responses and species divergence through time could weaken competition and promote coexistence.
2. Other seasons place stress on fish (e.g., summer can pose thermal or oxidative stress), but species diverge most strongly in their activity rates during winter. Some species are expected to be more successful foragers and competitors under the unique conditions of dark, cold, low oxygen, or low resource winter conditions than others. Fluctuation-dependent coexistence via temporal niche divergence could therefore arise most strongly during and around winter periods.
3. Trade-offs exist between performance in open water and winter. Species with a higher capacity for growth in warm or bright periods are less successful during winter periods and vice versa. Other potential trade-offs include thermal breadth and thermal preference (Fig. 2B), thermal breadth and maximum performance at optimal temperatures (Kingsolver 2009) and between maximum growth capacity and tolerance to harsh conditions (Angert et al. 2011).
4. The strength of competition and the extent of niche partitioning varies through time (among seasons and years) and space (among lakes).

a. Reduced availability or accessibility of resources occurs during winter and should amplify inter-specific competition compared to more productive summer months.
b. The strength of competition should be highest in small or less productive lakes and weakest in large or highly productive lakes.
c. However, competition could be readily dampened by resource or habitat partitioning or divergence of activity rates in a given season.
d. Warmer winters are predicted to have the largest negative effects on biodiversity in small or low productivity lakes. In such lakes, where the capacity for spatial resource partitioning is low, shortening winter periods could increase convergence and synchrony in annual activity patterns and amplify competition.
5. The importance of niche divergence in time vs. space in mediating coexistence is context dependent. Divergence through time could be most important, generally, in low productivity, small, or highly spatially homogenized ecosystems.
6. Alternatively, if species niche overlap does not respond to winter duration, then winter might be predicted to have minimal effects on biodiversity. If winter causes species to converge in their habitat use or activity level, then long winters might even have a negative effect on biodiversity. In such cases, shorter winters might have little or even positive effects on biodiversity.

## Conclusions

Temporal variation and periodicity are ubiquitous in nature. Temperate and Arctic latitudes are characterized by seasonal cycles with distinct periods of winter. Here, we have argued for a perspective that emphasizes the roles of seasonal variation and **winter-mediated species coexistence**. In lakes, winter creates a unique suite of environmental conditions, to which fishes have differentially adapted. Most fish are active and growing in the open water seasons, but only some species are efficient foragers and competitors in the dark and cold of winter. Competing species with different thermal preferences should predictably diverge in their capacity for winter activity (i.e., species with colder thermal preferences should be more winter active). Even species with similar thermal preferences that may both be capable of winter activity (e.g., if they both tolerate cold temperatures) could diverge in their performance during low light, oxygen, or resources under ice covered periods. These divergent responses to winter could play a widespread role in coexistence based on existing theory. New theory developed here further argues that: 1) the duration of winter could be critically important for facilitating coexistence in some instances (e.g., when competition during winter is strong), and 2) winter need not be ‘good’ for the growth of a winter-adapted species to promote their coexistence. Future work is tasked with exploring how the environmental context, including lake size, productivity, community, and food web structure (e.g. presence of particular prey, competitors and predators) governs the role of winter for shaping fish behavior, growth, and biotic interactions.

## Acknowledgements

BCM and MR thank NSERC Discovery Grants for funding and the Canada Research Chair program to MR.

## Author contributions

BCM conceived the idea for the manuscript and wrote the first draft. KSM conceived and developed the theory with CB. MMG, TB, PB, TF, HG, TM, MR, MR and BS contributed data and figures and all authors contributed to writing and editing and the manuscript.

