## Supplemental Information for "Winter in water: Differential responses and the maintenance of biodiversity"

**S1. Acoustic telemetry**

**S1.1. Lake of Two Rivers, Ontario, Canada**

*Telemetry Array*

The Lake of Two Rivers telemetry array consists of 54 Vemco 69 kHz VR2W omnidirectional receivers (Vemco Ltd., Bedford, NS, Canada) with co-located V16 sync tags (Fig. S1). The mean distance between receivers was 236 m.

Receivers were affixed at a depth of 2 m to vertical lines anchored to two 8 kg steel weights and suspended by a surface float. A synchronization tag also affixed to each line below the receiver. Floats were submerged beneath the lake surface in the fall to protect from the effects of ice over winter. Six reference tags (V9P) were deployed throughout the array at depths of 5 and 18 metres to provide fixed positions which could be used to evaluate the performance of the array.

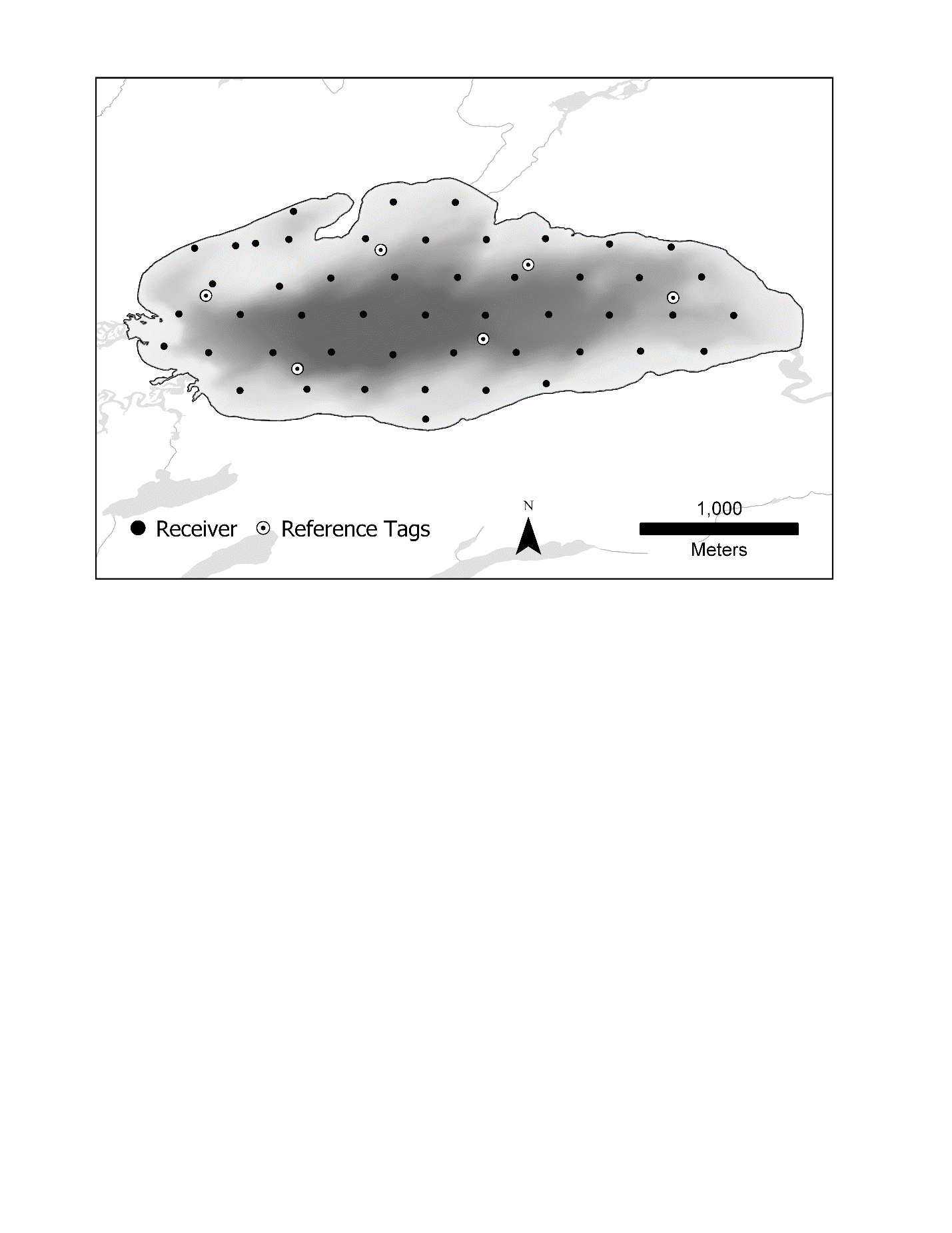
Two strings of temperature loggers (HOBO U22 Water Temp Pro, Onset Computer Corporation, Bourne, MA, USA) were also deployed, one at each end of the lake. Loggers were attached to the lines anchoring receivers at depths of 1.5, 3, 4.5, 6, 7, 8, 9, 10, 11.5, 13, 20, and 27 m in each location. Each logger was programmed to record temperature every 15 minutes.

**Figure S1.** Map of the telemetry array in Lake of Two Rivers. Black points indicate receiver and co-located sync tag positions. Open circles with black dots indicate fixed position reference tags.

*Tag implantation*

Surgeries to implant tags into lake trout and smallmouth bass were performed over three periods, May 2017, Sept. 2017, and May 2018. During each of these periods lake trout and smallmouth bass were captured either by angling, trapnet, or short set duration gillnet. Upon capture, if fish were deemed to be in good condition, they were anaesthetized using a buffered solution of tricaine methanesulfonate (MS-222), placed into a wetted foam surgery cradle where a flow of maintenance dose buffered MS-222 solution was directed across their gills. A small incision was made midline and posterior to the pectoral fins using a clean surgical scalpel and the telemetry tag was inserted. The incision was stitched with 2-3 sutures using 3-0 Monocryl synthetic absorbable sutures (Ethicon Inc, Somerville, NJ, USA). Length (±1 mm) and scale samples were obtained from the fish and an external t-tag was applied at the base of the dorsal fin. The total time for surgery and sampling was typically less than 3 three minutes per fish. The fish was then placed into a tank filled with lake water and allowed to recover until it was able to right itself and exhibited strong tailbeats, which typically occurred within 10-15 min. The fish was then returned to the lake.

Smallmouth bass and lake trout were implanted with pressure sensitive V9P (Vemco Ltd, Bedford, NS, Canada) tags programmed to transmit at a pseudo random interval, on average every 420 s, ranging between 360 and 480 s. Estimated tag life was 912 days. Table 1 provides details fish used in this analysis. The V9P tags were 9 mm X 31 mm and weighed 2.8 g in water.

**Table S1.** Lengths of fish tagged and used in this analysis

| Species | n | Mean Length (Min-Max) mm |
| --- | --- | --- |
| Lake trout | 17 | 424 (326 – 521) |
| Smallmouth bass | 15 | 382 (326 – 497) |

*Raw data processing*

Receivers were downloaded every six months. As the receiver retrieval, download, and redeployment process took several days, there are two intervals during the study period for which detection data are not used given the array was in a state of flux. Raw detection data were processed by Vemco using a hyperbolic positioning algorithm (Smith, 2013) to assign x,y positions to fish detected at multiple receivers over the course of the study resulting in 624,736 positions post processing between May 08, 2017 and October 30, 2018. Approximately 30% of these positions were from smallmouth bass, and 70% from lake trout.

*Array performance*

Fixed location reference tags within the array provided a measure of array performance throughout the study period. Estimated positions based on Vemco’s positioning algorithm were compared with known positions of the tags to quantify positioning error across seasons and years using methods proposed in (reference). Reference tags at 5 metres in depth had lower average error than tags placed at 18m with over 96% of detections having a positional error of less than 6m vs. 86% and an overall mean error of 2.25 metres vs 5.15 metres.

**Table S2.** Array performance at two depths of fixed tags. The three tags which were placed at each depth are pooled here for summary purposes.

| Tag Depth | Mean Error | Median Error | % of detections > 15m error | % of detections < 6m error |
| --- | --- | --- | --- | --- |
| 5 | 2.25 | 1.0 | 1.3 | 96.7 |
| 18 | 5.15 | 0.7 | 9.6 | 86.0 |

**S1.2. Alexie Lake, Northwest Territories, Canada**

Alexie Lake (62°40′36.59″ N, 114° 4′22.76″W) is a is a medium-sized (402 ha, maximum depth 32 m), oligotrophic lake located approximately 30 km north east of Yellowknife, Northwest Territories, Canada. Despite being located >60˚C latitude, Alexie Lake undergoes thermal stratification each summer months with peak surface water often exceeding 20˚C. More details on Alexie Lake can be found in (P. a. Cott, Johnston, & Gunn, 2011). Alexie Lake was outfitted with a temperature string and an acoustic array comprised of 72 omni-directional receivers from July 2012 through October 2014, inclusive. Details on the telemetry array, tags, and surgical information can be found in (P. A. Cott, Guzzo, Chapelsky, Milne, & Blanchfield, 2015; Guzzo, Blanchfield, Chapelsky, & Cott, 2016). For this study we used telemetry data for lake trout (n = 18) and burbot (n = 2) recorded between 27 June 2013 and 7 July 2014 (Table S3).

**S1.3. Telemetry Analysis (Lake of Two Rivers and Alexie Lake)**

*Analysis Filters*

The processed position data from Lake of Two Rivers and Alexie Lake were used to calculate daily average depth, daily average bathymetry position, and daily average movement rates of the two study species using all detections within the study periods defined above for each lake.

Positions recorded less than 14 days after tagging were excluded form the analysis to provide fish time to recover (Rogers and White 2007). In addition to this filter, specific filters were applied prior to calculating each of the metrics.

Prior to calculating average depth and bathymetry depth, detections were excluded if the recorded depth was more than 2 m deeper than the maximum lake depth. Recorded depths less than 0 metres were set to 0.

Prior to calculating average movement rate detections were excluded if the position fell outside the lake boundary, or the time between detections was more than 20 minutes. We then removed all speeds >40 m min^-1^, which represent impossible speeds. The final numbers of positions used in each analysis is provided in Table 3.

Spatial positions for each species during July and August 2017 (denoted summer) and January and February 2018 (denoted winter) were subset from the larger dataset and plotted out to help visualize the stark change in spatial habitat use by lake trout and smallmouth bass between summer and winter seasons (Fig. 3).

**Table S3.** Number of detections used per species from each telemetry study after data filtering.

| Lake | Species | Depth, Temperature,  Bathymetry Analysis | Movement Analysis |
| --- | --- | --- | --- |
| Lake of Two Rivers | Lake trout | 324,213 | 95,288 |
|  | Smallmouth bass | 249,401 | 84,627 |
| Alexie Lake | Lake trout | 1,155,805 | 59,332 |
|  | Burbot | 71,927 | 7,032 |

**S2. Fish growth**

**S2.1. Lake trout and small mouth bass, Lake Opeongo, Ontario**

*Individual fish growth measurements*

Growth was estimated from individual fish back-calculated length-at-age, so each individual could provide measurements for several years. The data included 214 lake trout and 879 small-mouth bass individuals, whose growth could be assessed from 1973 to 2005 and 1965 to 1988, respectively. More details of fish sampling and measurement methods can be found in Shuter et al. (1987, 2016). Winter phenology data were derived from measurements of freeze-up and ice break-up days in Sproules Bay within Lake Opeongo by the Ontario Ministry of Natural Resources and Forestry. Freeze-up days have been measured since 1964, and break-up day data range from 1978 to 1999. A schematic summary of the winter phenology variables and the associated growth variable is depicted in Figure S2. They were chosen to represent any immediate or adjacent effect of phenology on growth, so they include both the previous winter and winter following the target growing season.

Growth (cm/year) was calculated for each fish ‘i’ and year ‘y’ combination as:

$${Growth}_{i,y}=L_{i,y}-L_{i,y-1} (S.1)$$

Where *L* is fork length (cm). Year ‘y’ represents “current” year, where most of the growth is expected to happen (i.e., it includes the summer), so $L_{i,y}$ is the length at the end of year ‘y’, and $L_{i,y-1}$ is the length at the end of previous year (Figure S1).

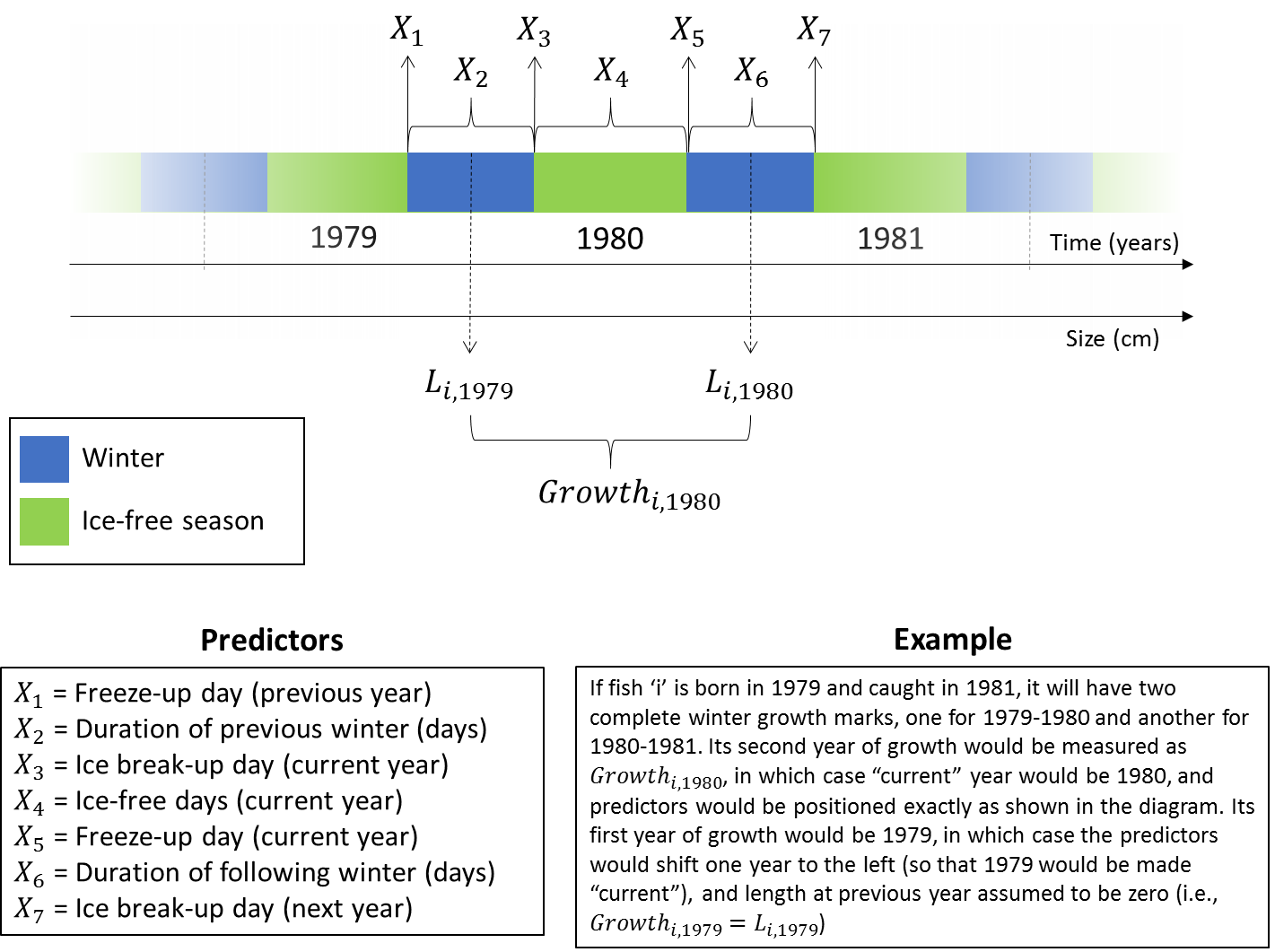

**Figure S2.** Winter phenology variables used to predict growth rates of individual lake trout and small mouth bass in Lake Opeongo.

*Statistical analyses*

Growth data were analyzed through Generalized Linear Mixed-effects Models (GLMMs) in MATLAB R2018b. Each predictor X was included in a separate model, coded as:

$$Relative Growth \sim1+X*Species+Age*Species+\left( 1+Age | Fish \right) (S.2)$$

The response variable, relative growth, is the individual growth divided by the species’ maximum length. It was chosen as the response variable to standardize for differences in species overall length and shape (i.e., small-mouth bass has a deeper body, so growth in fork length is expected to be slower in absolute terms, although it is the relative effect of winter phenology variables that we are interested in). The maximum length was calculated as the mean fork length of the ten largest fish in the sample, as in McDermid et al. (2010), resulting in 85cm for Lake Trout and 42cm for Small-mouth Bass. The model includes the main effects of X, Age, and Species, as well as the two-way interactions depicted in Equation S.2. We were particularly interested in the predicted slopes of X for each species, which can be calculated for lake trout (used as reference species) as the main effect of X and for small-mouth bass as the main effect plus the interaction X*Species. Individual Fish was used as a random factor, with random variation in both intercept (overall growth) and slope (how growth changes with Age).

Only immature growth was considered for analysis. For Lake Trout, age at maturity was defined individually, according to Shuter et al. (2016). For Bass, growth was calculated up to 4 years of age, based on conservative estimates of age at maturity in the lake (Dunlop et al. 2005a,b). Even after controlling for potential effects of maturation and reproductive costs, growth trajectories could still be non-linear, and for this reason Age was included as a covariate. For the first year of growth, the initial length was assumed as zero, i.e., $L_{i,0}=0$. The results remain qualitatively the same if this first year is excluded from analysis.

The predicted slopes show a much more pronounced response of small-mouth bass to changes in winter phenology then lake trout (Table S4). The signs of slopes are consistent with slower small-mouth bass growth in years that were preceded by longer winters (earlier freeze-up day X_1_, later ice break-up day X_3_ and consequently longer winter duration X_2_). Figure 4A,B in the main text shows the partial residual plots for the model using X_2_ as a predictor. Another interesting result is that the following winter had the opposite effect on small-mouth bass growth: earlier freeze-up day X_5_, later ice break-up day X_7_ and longer winter duration X_6_ were associated with faster relative growth. This can be interpreted as a by-product of size-selective winter mortality (e.g., Shuter and Post 1990), i.e., only fish that had grown fast enough that year would be able to survive through the next winter and be captured in one of the following years. These two opposing effects might explain why the number of ice-free days (X_4_) had no effect on growth.

**Table S4.** Statistics from GLMMs, each using a winter phenology variable identified by the “Predictor” column. n = number of observations (fishes and ages per fish) used to fit the model; R^2^ = adjusted R^2^; LT = lake trout, SMB = small-mouth bass. Slopes represent the predicted fixed effects of the predictor variable on relative growth of each species, those highlighted in bold are different from zero (p<0.05). The reference ages used to calculate the slopes were 4.6 and 2.5 years for lake trout and small-mouth bass, respectively.

| Predictor | n | R^2^ | LT slope x 10^4^ | SMB slope x 10^4^ |
| --- | --- | --- | --- | --- |
| X_1_: Freeze-up day (previous year) | 2390 | 0.58 | -0.506 | **18.172** |
| X_2_: Duration of previous winter | 2390 | 0.60 | -0.087 | **-15.456** |
| X_3_: Ice break-up day (current year) | 5235 | 0.52 | -0.313 | **-1.417** |
| X_4_: Ice-free days (current year) | 2319 | 0.56 | 0.167 | -1.708 |
| X_5_: Freeze-up day (current year) | 2319 | 0.57 | 0.002 | **-9.946** |
| X_6_: Duration of following winter | 2319 | 0.57 | 0.087 | **8.162** |
| X_7_: Ice break-up day (next year) | 5235 | 0.52 | 0.061 | **4.654** |

**S2.2. Lake trout, IISD-Experimental Lakes Area**

Lake trout growth was examined using data from annual mark–recapture sampling data from two lakes (Lakes 224 and 373) within the IISD-Experimental Lakes Area (IISD-ELA). Fish were captured each fall using trap nets and short (<30 min) evening gill net sets on spawning shoals (Mills et al. 1987, 2002). Following capture, the weights, fork lengths (in millimeters), and tag numbers from previously captured fish were recorded. Newly captured fish received a tag for future identification. We identified 241 and 1295 instances in which an individual fish was captured in consecutive years during the study period, in Lake 373 and Lake 224, respectively. We then related the change in mass (g) between these two captures to the duration of ice-cover that fell between the two capture periods. Annual duration of ice-cover for the study lakes was inferred from ice-formation and ice-breakup dates from Lake 239 (Rawson Lake), also within the IISD-ELA. Since the lakes are all similar size and are located within 10 km from one another we assumed ice-cover was similar across the lakes (Guzzo & Blanchfield, 2017; Guzzo, Blanchfield, & Rennie, 2017).

Linear regression results for change in annual body mass as a function of ice-cover duration.

Lake 373: est = -0.68 ± 0.11, t = -6.08, p < 0.001

Lake 224: est = -0.73 ± 0.30, t = -2.43, p = 0.01

**S3. Fish Bioenergetics**

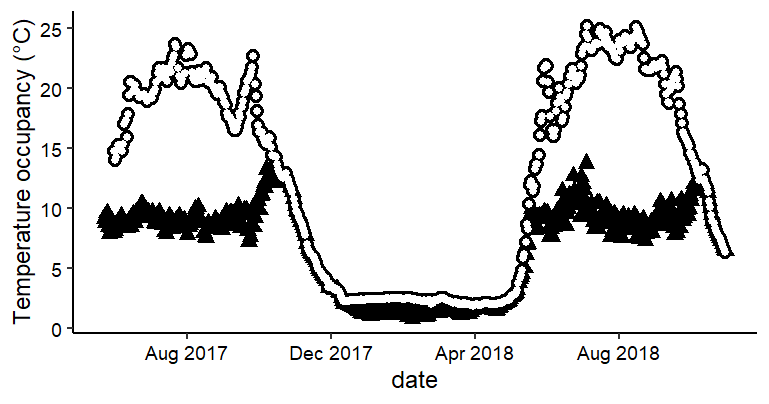
We used the mean daily temperature occupancy values obtained for lake trout and smallmouth bass from telemetry in Lake of Two Rivers, Ontario to estimate the daily consumption estimates which these fish would need to maintain their weight over the course of one year using the Wisconsin Bioenergetics Model (Hanson, Johnson, Schindler, & Kitchell, 1997). Models parameterized for each species were run in the R package Fish Bioenergetics 4.0 (FB4) (Deslauriers, Chipps, Breck, Rice, & Madenjian, 2017). Mean daily temperature occupancy for each species estimated were obtained by translating each depth detection to the mean daily water temperature recorded at that depth from the temperature string (Fig. S2). Starting mass and energy density for each species was 400 g and 5510 J/g. Diet of each species was the same with a set energy density of 3500 J/g (10% indigestible). Because simulations used of the same start and end mass (400 g), we were modelling how much food each species would have to consume to maintain their starting mass over the course of one year, given the average water temperatures each species was exposed to. The model also estimates the corresponding daily respiration rates.

**Figure S3.** Mean daily temperature occupancy of lake trout (black triangles) and smallmouth bass (white circles) in Lake of Two Rivers, Ontario recorded during 23 May 2017 and 30 October 2018.

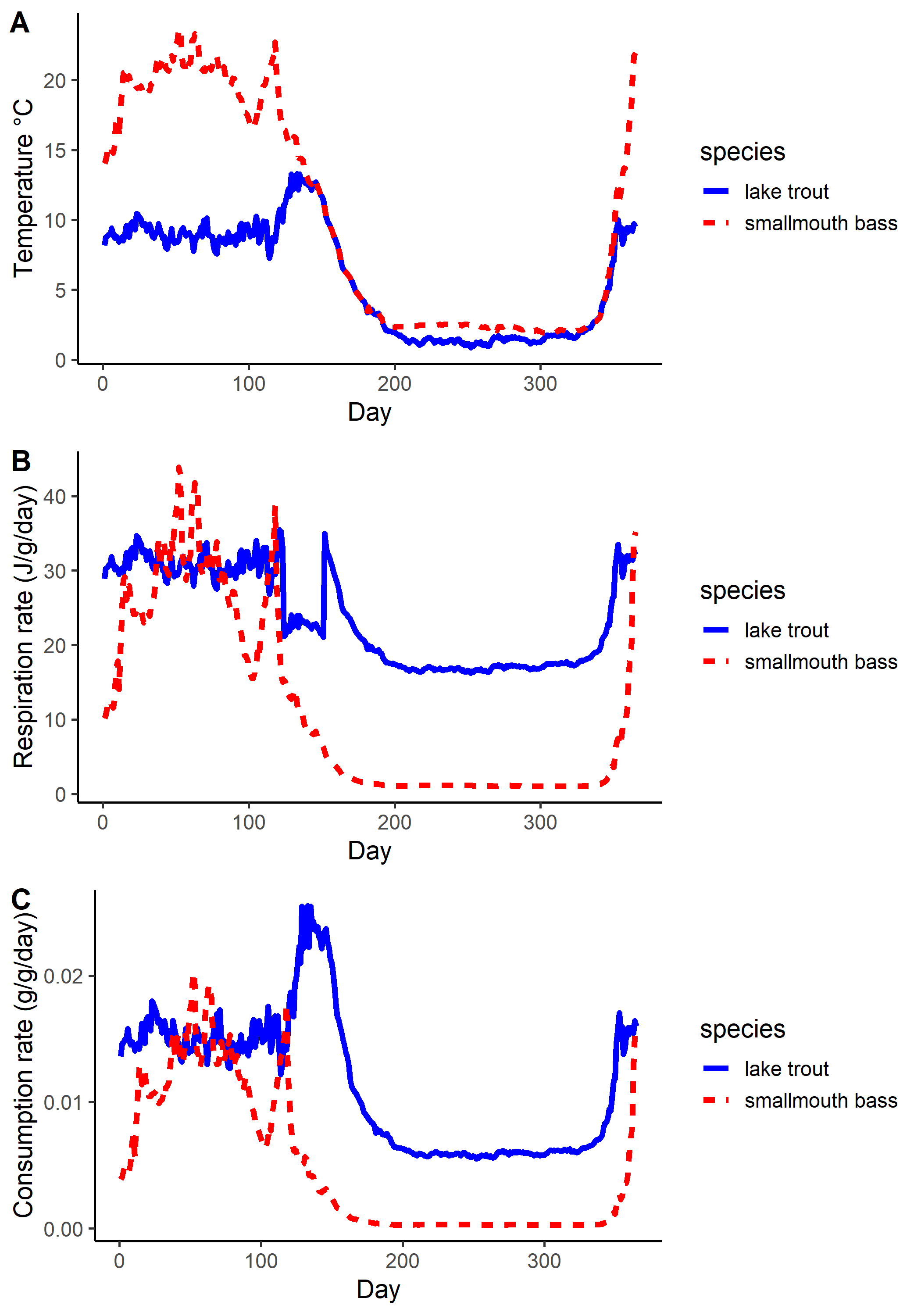

**Figure S4.** Output of bioenergetic model simulations for lake trout and smallmouth bass to maintain their starting weight of 400 g over the course of one year based on temperature occupancies from telemetry data collected in Lake of Two Rivers, Ontario (Day 1 = 1 June 2017 through Day 365 = 30 May 2018). A. Mean daily temperature occupancy of each species during the simulation. B and C Modelled daily specific respiration rate (J/g/day) and consumption rates (g/g/day).

**S4. Theory**

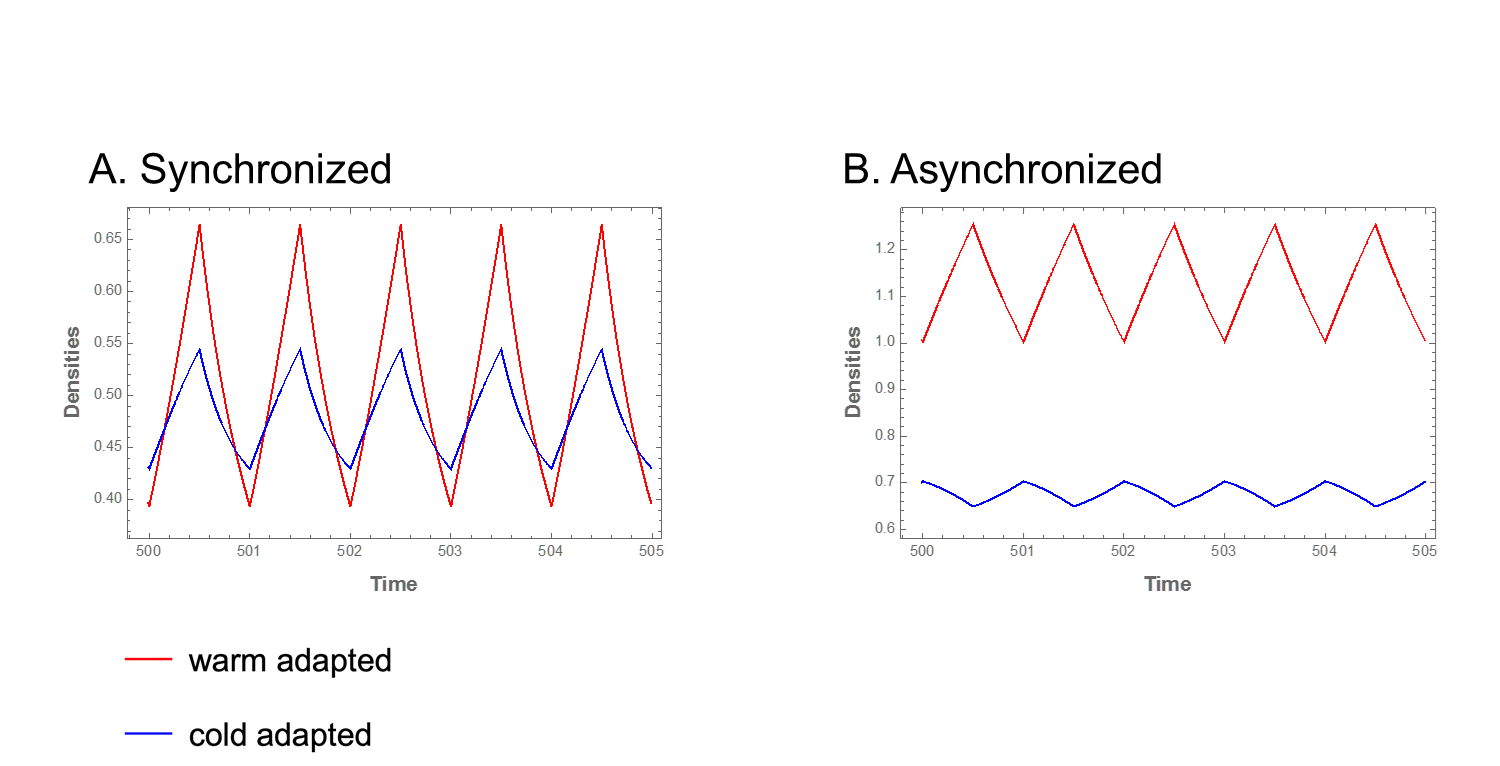

**Fig. S5** Time series examples for the strong-strong case showing that dynamics can be either synchronized such that both species decline in winter, or asynchronized such that the cold-adapted species increases during the winter. Importantly, parameters used to generate both time-series patterns for all of the above 4 cases produce qualitatively similar outcomes for coexistence.
